# Role of enhanced glucocorticoid receptor sensitivity in inflammation in PTSD: Insights from a computational model for circadian-neuroendocrine-immune interactions

**DOI:** 10.1101/664201

**Authors:** Pramod R. Somvanshi, Synthia H. Mellon, Rachel Yehuda, Janine D. Flory, Linda Bierer, Iouri Makotkine, Charles Marmar, Marti Jett, Francis J. Doyle

## Abstract

Although glucocorticoid resistance contributes to increased inflammation, individuals with post-traumatic stress disorder (PTSD) exhibit increased glucocorticoid receptor (GR) sensitivity along with increased inflammation. It is not clear how inflammation co-exists with a hyper-responsive hypothalamic pituitary adrenal axis (HPA axis). To understand this better, we developed and analyzed an integrated mathematical model for the HPA axis and the immune system. We performed mathematical simulations for a dexamethasone suppression test and IC_50_-dexamethasone for cytokine suppression, by varying model parameters. The model analysis suggests that increasing the steepness of the dose response curve for GR activity may reduce anti-inflammatory effects of GRs at the ambient glucocorticoid levels thereby increasing pro-inflammatory response. The adaptive response of pro-inflammatory cytokine mediated stimulatory effects on the HPA-axis is reduced due to dominance of the GR-mediated negative feedback on the HPA-axis. To verify these hypotheses we analyzed the clinical data on neuro-endocrine variables and cytokines obtained from war-zone veterans with and without PTSD. We observed significant group differences for cortisol and ACTH suppression tests, pro-inflammatory cytokines TNFα and IL6, hs-CRP, promoter methylation of GR gene and IC_50_-Dex for lysozyme suppression. Causal inference modelling revealed significant associations between cortisol suppression and post-dex cortisol decline, promoter methylation of NR3C1-1F, IC_50_-Dex and pro-inflammatory cytokines. We noted significant mediation effects of NR3C1-1F promoter methylation on inflammatory cytokines through changes in GR sensitivity. Our findings suggest that increased GR sensitivity may contribute to increased inflammation, therefore, interventions to restore GR sensitivity may normalize inflammation in PTSD.

## Introduction

The HPA axis is one of the major regulatory pathways that controls the physiological response to stress. Alterations in the response of GRs in HPA axis are implicated in the pathogenesis of PTSD (Yehuda, 2006). Since the HPA axis also regulates inflammatory responses, changes in HPA-axis activity are often associated with increased inflammation as observed in several psychiatric disorders. Glucocorticoids are the mediators of the HPA axis that operates through glucocorticoid receptors to exert their action on physiological functions. Studies have reported varying findings on glucocorticoid receptor (GR) number, GR promoter methylation and GR mRNA expression levels in PTSD (Geuze et al., 2012; Gotovac et al., 2003; Labonté et al., 2014; Liberzon et al., 1999; Logue et al., 2015; Martins et al., 2017; Rachel Yehuda, 1991; van Zuiden et al., 2012; Vukojevic et al., 2014; Yehuda et al., 1995; Yehuda et al., 1993; Yehuda et al., 2015). PTSD studies have reported inconclusive findings in regards to cortisol level and GR sensitivity (de Kloet et al., 2007; Lindley et al., 2004; Matić et al., 2013; Shalev et al., 2008; Wheler et al., 2006; Yehuda, 2009; Yehuda et al., 1995). However, findings supporting the enhanced sensitivity of GR and increased inflammation are consistent across the majority of PTSD cohorts (de Kloet et al., 2007; de Kloet et al., 2006; Gill et al., 2010; Lindqvist et al., 2017; Rohleder et al., 2004; Rohleder et al., 2010; Vidović et al., 2007; von Känel et al., 2007).

Contrary to other psychiatric disorders that exhibit increased plasma levels of both cortisol and pro-inflammatory cytokines suggesting possible glucocorticoid (GC) resistance, PTSD is associated with enhanced GR sensitivity and inflammation (Hoge et al., 2009; Neigh and Ali, 2016; Wang et al., 2017). Although GC are well known for their anti-inflammatory properties, there are also accounts of the pro-inflammatory effects of GCs (Cruz-Topete and Cidlowski, 2015; Desmet and De Bosscher, 2017), where low dose glucocorticoids are associated with enhanced inflammatory responses and high doses are anti-inflammatory. It is unclear how GCs exhibit both anti-inflammatory and pro-inflammatory effects. It is hypothesized that the decreased cortisol levels, as observed in the majority of PTSD subjects, may reduce the immunosuppressive effects of cortisol thereby increasing inflammation. On the other hand, there are accounts of increased cortisol levels as well as increased inflammation, as observed in some PTSD subjects and other psychiatric disorders with GC resistance. Since there is not a unidirectional relationship between cortisol and inflammatory cytokines and because the HPA-immune axis is composed of multiple feedback loops mediated by glucocorticoid receptors and cytokines, it is important to analyze these relationships from a systems perspective (Newton et al., 2017).

The HPA axis is a homeostatic system composed of CRH, ACTH and cortisol. Stress activates CRH in the hypothalamus that influences cortisol secretion from adrenal cortex through activation of ACTH in pituitary. Cortisol negatively regulates CRH and ACTH production through binding to GRs. GRs are expressed in several brain and peripheral tissues and regulate neural, metabolic and inflammatory responses to stress on binding of cortisol (Oakley and Cidlowski, 2013). While GCs show nonlinear dose-response effects on pro-inflammatory cytokine (TNFα and IL6) expression (enhanced at low doses and inhibited at higher doses), cytokines are known to activate the HPA axis at all three levels, and inhibit GR nuclear activity (Van Bogaert et al., 2010; Webster et al., 2001). Therefore, the HPA axis can also be activated by inflammatory stimuli such as LPS or infection. The interactions between inflammatory signaling and the HPA axis are composed of multiple feedback loops including crosstalk between the two pathways (Cain and Cidlowski, 2017) (See Figure 2A).

GRs are expressed in peripheral tissues and lymphocytes, and its signaling cascade is initiated by binding of circulating glucocorticoids (Yehuda and Seckl, 2011). High doses of glucocorticoids typically inhibit pro-inflammatory cytokine expression and activity (Brattsand and Linden, 1996; Coutinho and Chapman, 2011). Whereas, the pro-inflammatory cytokines (TNFα and IL6) are known to activate the HPA axis by their direct and indirect action on CRH, ACTH and cortisol secretion (Turnbull and Rivier, 1999). These cytokine-HPA-axis interactions can be viewed as an actuator-dependent secondary negative feedback that acts by reducing a positive effect of cytokine on cortisol release. GR nuclear action also negatively feeds back on its mRNA synthesis. At the systems level, these multiple feedbacks operate together to ensure homeostasis. The state of the glucocorticoids and the inflammatory cytokines are determined by the strength and functioning of these feedback loops and the crosstalk between the two pathways. Therefore to understand the variability in the response of HPA-immune axis we developed and analyzed a mathematical model of the HPA-Immune axis.

To characterize the HPA-immune axis relationships, we obtained data for cortisol, ACTH, dexamethasone suppression test for cortisol and ACTH, IC_50_-DEX, methylation of the GR receptor gene and serum cytokine concentrations (IL6, TNFα, IFNγ, and IL10) and hs-CRP from 162 combat trauma exposed veterans, half of whom developed PTSD and the other half served as controls. We have reported earlier on significant associations between measures of GR sensitivity (dexamethasone suppression test and IC_50_) and both cortisol decline and methylation of the NR3C1-1F promoter in a subsample of the current cohort (n=59/59, case/control) and in other PTSD samples (Yehuda et al., 2015; Yehuda et al., 2003) and have separately reported an increased pro-inflammatory markers in our cohort (Lindqvist et al., 2017; Lindqvist et al., 2014) and stress induced inflammation in mouse models (Muhie et al., 2017).

In the current study, we studied the relationship among neuro-endocrine measures and cytokine assays and delineated the mechanism by which the HPA-axis measures may be related to an increased inflammatory response in PTSD using simulations of a mathematical model. We further verified the model-based mechanistic hypothesis through correlational analysis and causal inference using our clinical data. We developed and analyzed a systems level circadian model integrating the HPA axis with the inflammatory pathway parameterized on human data from the literature. We used model simulation to reconcile the discrepant findings of plasma cortisol levels, GR expression and IC_50_-DEX lysozyme suppression from different studies on PTSD cohorts reported in the literature. Based on model analyses, we propose a plausible mechanism for variability of HPA-axis and inflammatory responses as observed from group differences in features of these pathways in our data and other data reported in literature for PTSD subjects. We also explored whether changes in methylation of the NR3C1-1F promoter could be responsible for increased GR sensitivity and subsequent inflammation, in our sample using mediation analysis. Our analysis suggests that increased GC sensitivity contributes to enhanced inflammation in PTSD (Zhu et al., 2017), and the lower methylation of NR3C1-1F promoter contributes to increased GR sensitivity possibly through increased GR availability/expression.

## Results

### Group differences in HPA-axis and immune markers

Our sample was composed of combat exposed individuals who served in Iraq and Afghanistan, half of which had PTSD (n=81) and half did not (n=81, controls). The demographics our sample are reported in supplementary Table S1. Group means, standard deviations, fold change, p and q values are reported in Table S2. The boxplots for the HPA-immune variables are shown in Figure 1. We observed statistically significant differences in cortisol (p=0.058, n=162), ACTH (p=0.033, n=160), cortisol suppression (p=0.037, n=152), ACTH suppression (p=0.017, n=152), extent of cortisol decline (p=0.008, n=152) and ACTH decline (p=0.005, n=152) in response to dexamethasone. We also noted significant differences in IL6 (p=3.21E-4, n=160), TNFα (p=0.007, n=162) and hs-CRP (p=0.007, n=158); and a trend level difference in IC_50_-Dex (p=0.048, n=159) for lysozyme expression, methylation of the NR3C1-1F promoter region (p=0.027 adjusted for non-converted cytosine, n=161) and IFNγ. While cortisol, ACTH, cortisol and ACTH suppression, IL6, TNFα and hs-CRP were increased; IC_50_-DEX and GR promoter methylation were reduced in the PTSD subjects compared to controls. These overall indicated enhanced HPA activity, increased pro-inflammatory response along with increased central and immune sensitivity of glucocorticoid receptors in PTSD subjects.

**Figure 1.**
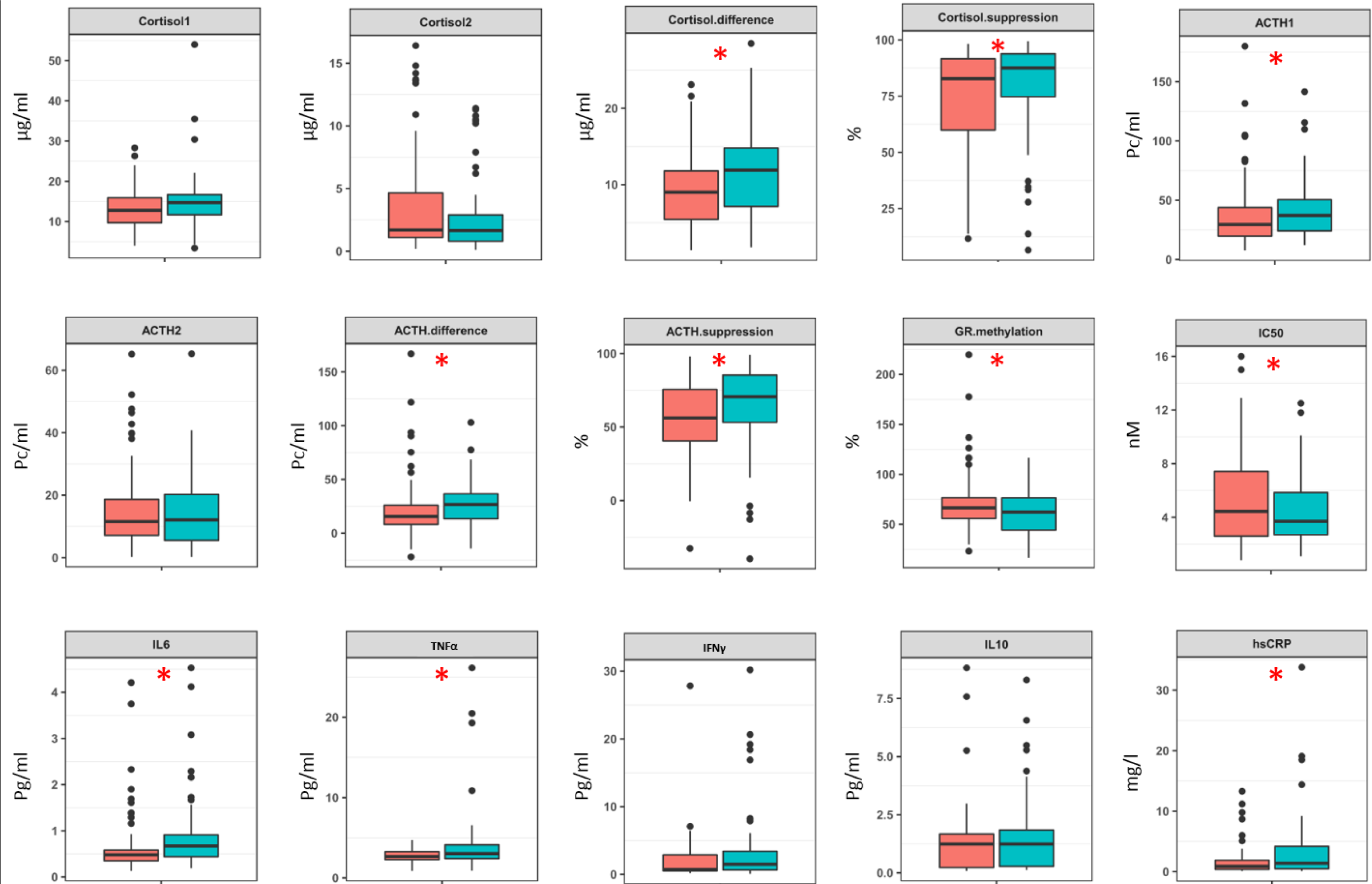
Boxplot representation of the group differences between the features of the HPA axis and cytokines in controls and PTSD subjects. The red and green box represents controls and PTSD data, respectively. The group means for cortisol difference, cortisol suppression, ACTH1, ACTH difference, ACTH suppression, IL6, TNFα and hs-CRP are significantly different with a trend in cortisol1 (pre-dex cortisol). The red star indicates the p<0.05. The p-values for the features are reported in Table S2.

**Figure 2.**
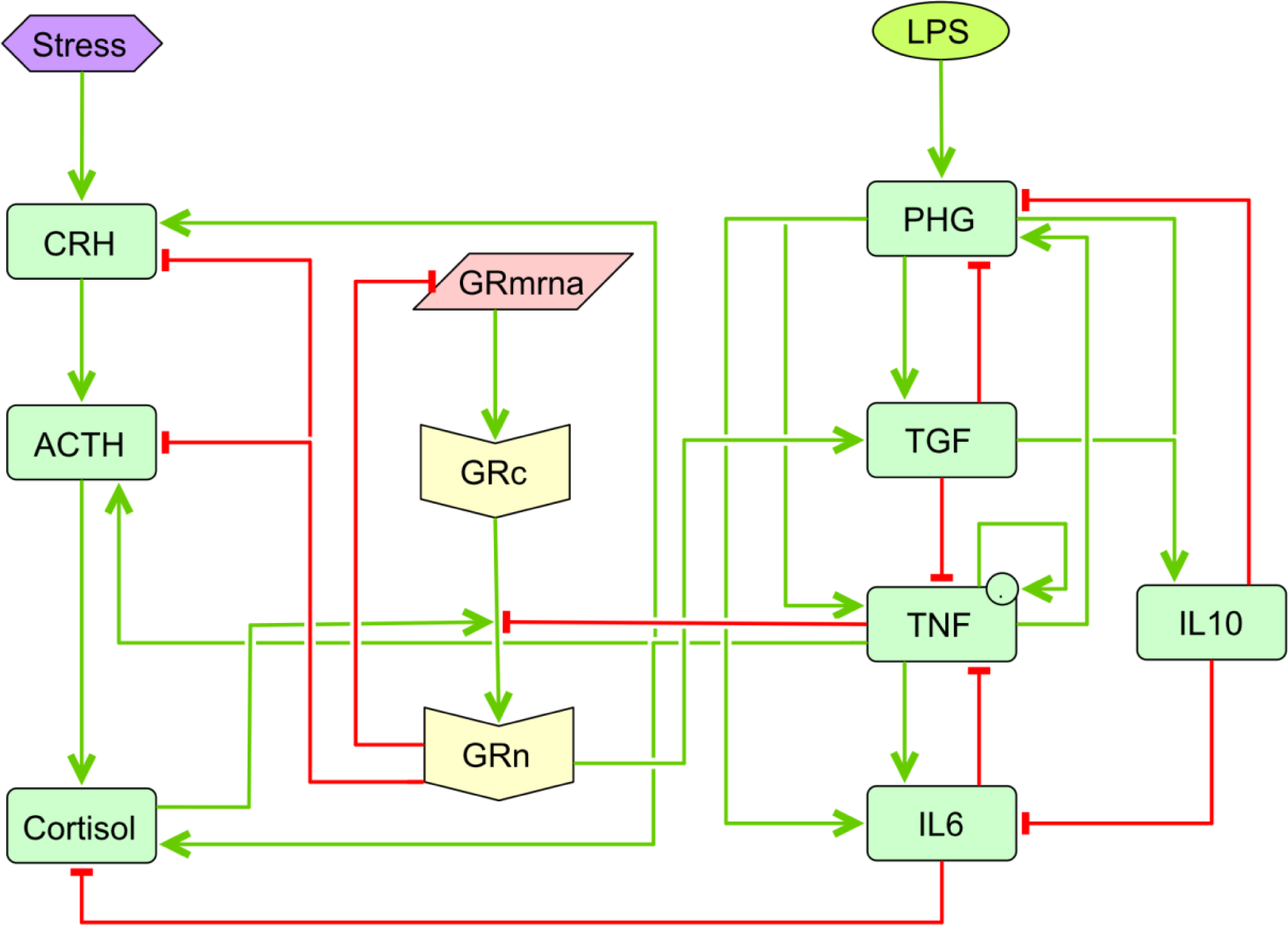
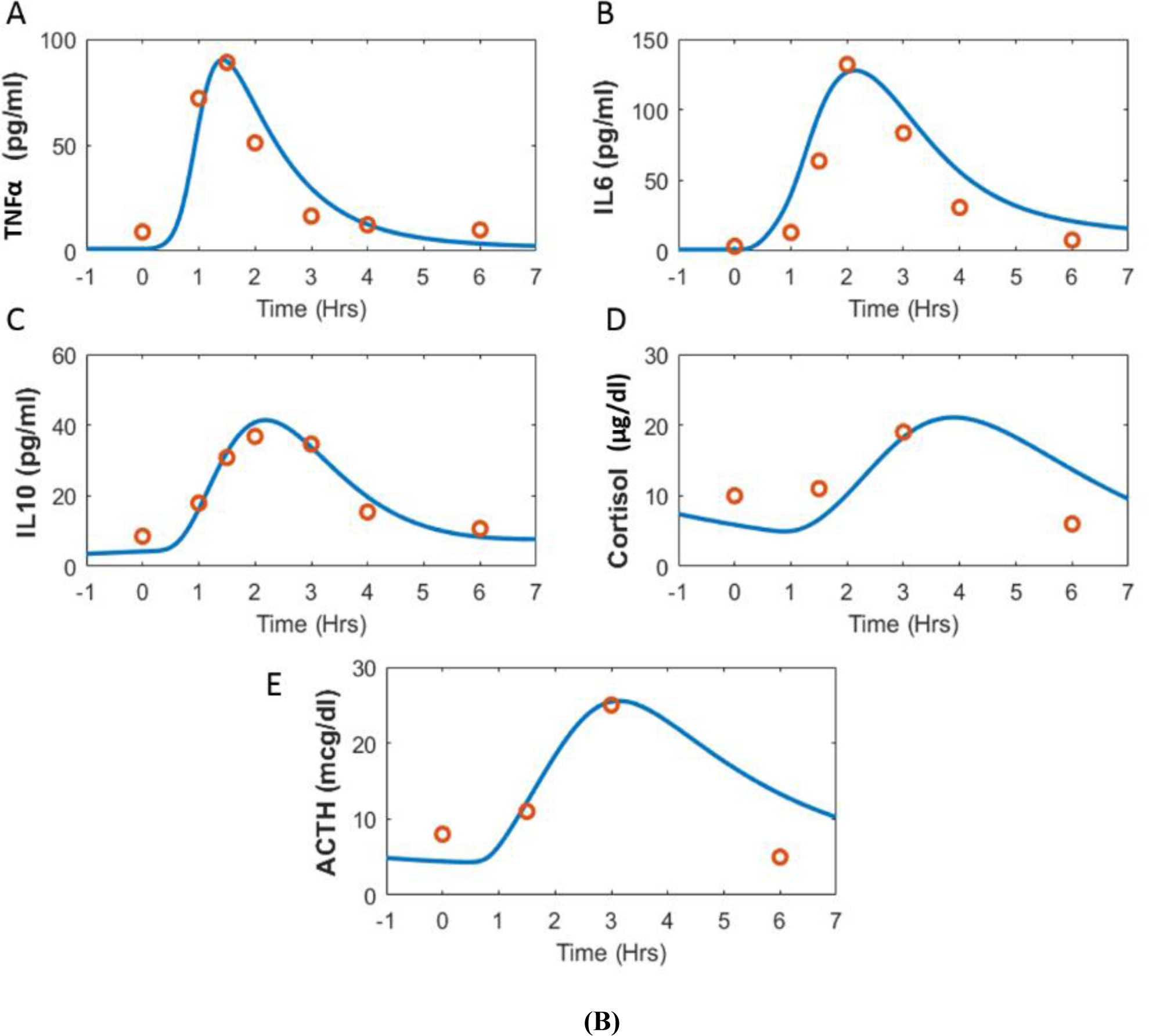
(A) The network representing the interaction between the HPA axis and the inflammatory pathway. The network is composed of multiple feedback loops and crosstalk between the two pathways. Glucocorticoid receptor dynamics lie central to the interactions between the two networks. Stress stimulates cortisol production though production of CRH and ACTH. Green arrows represent stimulatory effects and red arrows represent inhibitory effects. Cortisol increases GR nuclear translocation which negatively feedback on the upstream pathway. LPS activates phagocytes which in turn increase expression of both pro and anti-inflammatory cytokines. Anti-inflammatory cytokines inhibit phagocytosis and pro-inflammatory cytokines to restore the inflammatory response. Pro-inflammatory cytokines activate the HPA axis at CRH, ACTH and cortisol synthesis levels and inhibit GR nuclear translocation. Glucocorticoids in turn suppress inflammatory response through GRs. GRs are regulated through auto-positive and auto-negative feedbacks at its mRNA and protein synthesis. These multiple feedback mechanisms orchestrates together to restore homeostasis under the conditions of stress and inflammatory stimulus. (B) The simulated profiles of the cytokines and the HPA-axis variables for endotoxin administration at 0.4 ng/kg LPS dose and cross-validated against the clinical data (red circles) reported in (Grigoleit et al., 2010). The LPS dose was introduced at 0 Hrs. on X-axis that represents afternoon circadian time to mimic usual experimental conditions.

### HPA-Immune axis and mathematical model

We developed an integrated mathematical model of the HPA-immune axis network based on biological mechanisms reported in the literature and asked if it could predict biological observations on enhanced glucocorticoid sensitivity and inflammation previously reported in our studies. We adopted an integrated model structure for the HPA-axis and immune system interactions from the literature (Bangsgaard., 2016; Parker et al., 2016; Rao et al., 2016; Spiga et al., 2017), and updated the additional interactions of cytokines with the HPA axis. We parameterized the model for the human LPS stimulation response based on data reported in the literature (Copeland et al., 2005; Lauw et al., 2000; Wegner et al., 2017). We also incorporated the model for the pharmacokinetic-pharmacodynamics (PK-PD) response of dexamethasone for human data (Loew et al., 1986; Queckenberg et al., 2011; Spoorenberg et al., 2014). The validations of the model response of cortisol, ACTH, TNFα, IL6 and IL10 for a 0.4 ng/kg dose of LPS as reported in literature (Grigoleit et al., 2010) are shown in Figure 2B. Moreover, the model includes circadian dynamics of the HPA axis with two separate circadian drives representing the circadian drive from the suprachiasmatic nucleus (SCN) that regulates CRH release and the circadian drive from the adrenal clock that regulates StAR protein synthesis. The circadian drives were calibrated to attain the circadian profiles of the HPA axis components with an observed phase difference with reference to Cry1 of ∼6hrs between SCN and adrenal clocks (Pett et al., 2018). The detailed parameters are reported in Supplementary file Table S3.

### Model equations for integrated HPA axis-inflammation model

Corticotrophin releasing factor

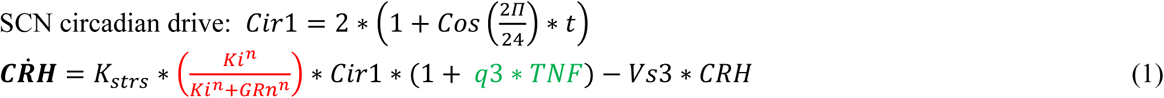

Adrenocorticotropic hormone

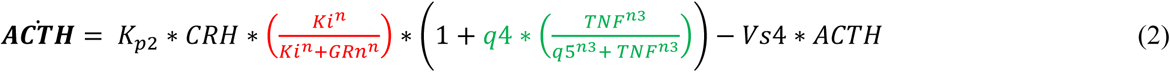

STAR protein

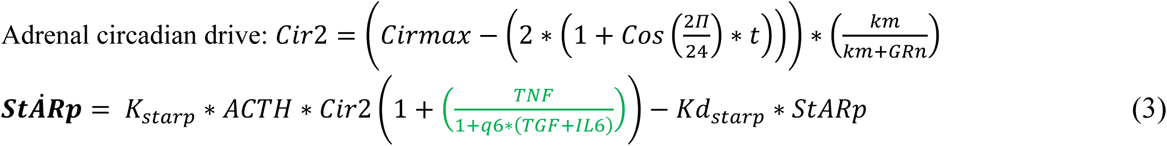

Plasma cortisol

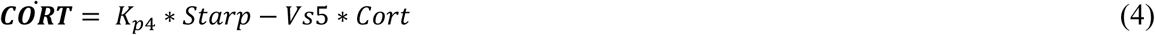

Dexamethasone compartment 1

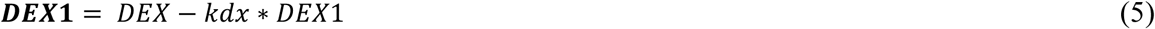

Dexamethasone plasma

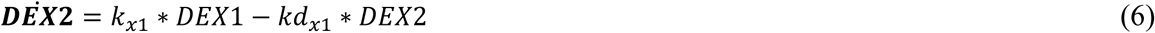

Delayed cortisol action

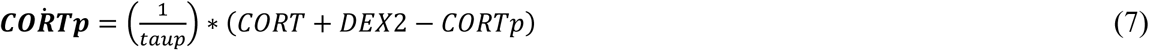

Glucocorticoid receptor mRNA

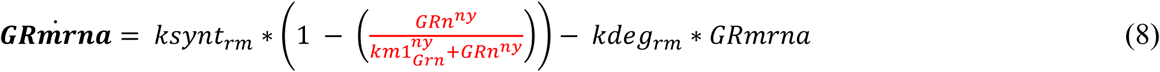

Glucocorticoid receptor protein

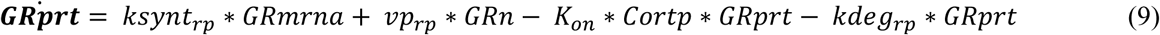

Cytosolic glucocorticoid receptors

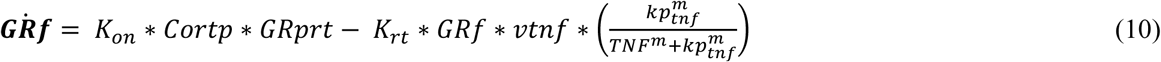

Nuclear glucocorticoid receptor

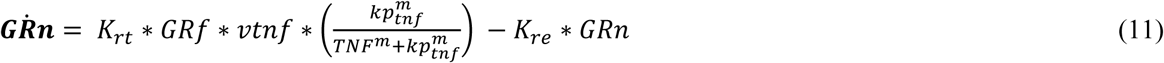

Lipopolysaccharides

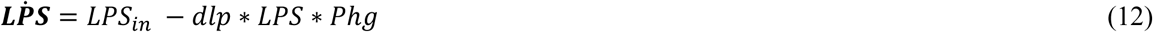

Phagocytes

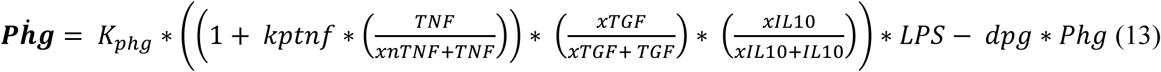

Transforming growth factor cytokine

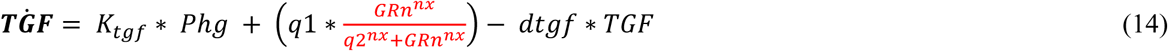

Tumor necrosis factor cytokine

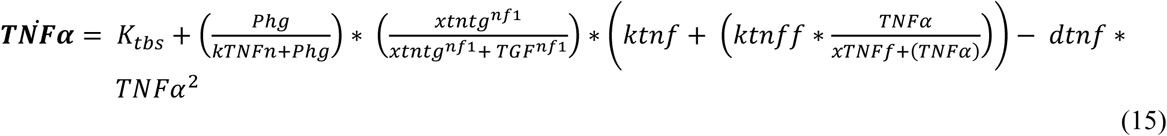

Interleukin 10 cytokine

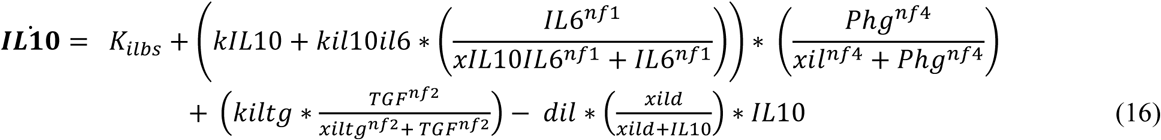

Interleukin 6 cytokine

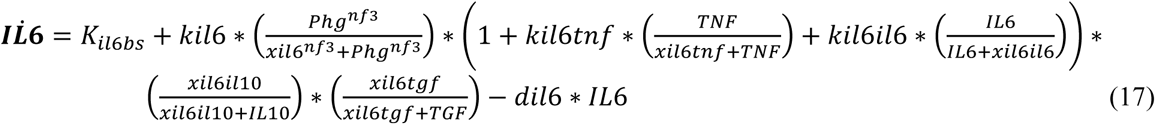

The expressions highlighted in red in equation no 1, 2, 8 and 14 represent the GR mediated regulation of the HPA-immune axis. The expression in Eqn. 1 and 2 represent negative feedback of GR on CRH and ACTH, respectively. The expression in Eqn. 8 represents the GR mediated auto-regulatory negative feedback on GR mRNA synthesis (Okret et al., 1986). The glucocorticoid mediated anti-inflammatory effect through activation of GR signaling is represented in expression in Eqn.14 (Coutinho and Chapman, 2011). The influence of pro-inflammatory cytokines on the activation of HPA-axis (Dunn, 2000) are represented by the expressions highlighted in green in Eqn. 1, 2 and 3. The distinct effects of the total regulation of the HPA axis is further explored in this model.

At the molecular level, the changes in GR sensitivity could be determined by several factors such as, intracellular hormone availability, GR expression levels, receptor isoform expression, hormone binding affinity, conformational changes due to hsp-complex, GR phosphorylation, nuclear translocation, DNA/GRE binding, interaction with other nuclear factors and circadian dynamics (Bamberger et al., 1996; Binder, 2009; Nicolaides and Charmandari, 2017; Nicolaides et al., 2014). The changes in any of these factors affect the steepness of the dose response curve of the GR activity. Therefore, in the current analysis, we varied the Hill coefficients, the measure of steepness in the dose-response effect in the expression highlighted in the red in the above equations to simulate the changes in GR sensitivity.

### Model simulations for cortisol suppression test

The group differences in our data suggest the coexistence of increased features of HPA-axis, GR sensitivity, and inflammatory response. To analyze the plausible mechanisms behind these observations, we performed simulations using the mathematical model described previously. A decline in cortisol following dexamethasone ingestion is greater in individuals with PTSD, suggesting decreased cortisol production due to increased GR sensitivity (Leistner and Menke, 2018; Yehuda and Seckl, 2011) Therefore, we simulated the dexamethasone suppression test for varying GR sensitivity in the model and recorded corresponding changes in cortisol levels, cytosolic GR receptor levels and inflammatory cytokines TNFα and IL6 levels

We performed simulations for varying systemic GR sensitivity, varying the sensitivity of both HPA axis and inflammatory pathways at the same time, with an assumption of consistent change in sensitivity across brain and immune cells. In mathematical terms, sensitivity of a dose response curve is denoted by the Hill coefficient in the biochemical kinetics (Somvanshi and Venkatesh, 2013), therefore the sensitivity of GR’s regulatory effect that captures the steepness of the response of GR’s downstream effects was modelled accordingly. In the model, the parameters for the equations representing the GR feedback loop were varied for simulating the changes in GR sensitivity. Each of these expressions have their respective Hill coefficient (sensitivity parameter) and the saturation thresholds (inhibitory or stimulatory) that is an amplification parameter. In these equations the Hill coefficients were simultaneously varied from 50%, 2-fold and 2.5-fold increase above the normal parameter values.

The dexamethasone induced cortisol suppression test is used to access the sensitivity of the GR negative feedback in the HPA-axis. We simulated this test using the mathematical model. The 0.5mg dexamethasone dose was introduced at 23:00 circadian time and the subsequent cortisol levels at 08:00 circadian time were recorded to evaluate post-DEX cortisol suppression. The simulation results are shown in Figure 3A. The pre-dexamethasone (pre-DEX or cortisol1) difference in levels of the variables represent the profiles for the effect of change in GR sensitivity. The dotted curve represents the normal profiles for native sensitivity levels. It can be noted that with increasing sensitivity, the cortisol and GR levels vary nonlinearly; wherein, for a modest 50% increase in sensitivity, the cortisol levels decreased and GR levels increased. For a 2-fold rise in sensitivity, the cortisol and GR levels showed little change. On further increasing sensitivity by 2.5 fold, cortisol levels rise and GR levels decrease. However, the cortisol suppression (CS) increased monotonically with increasing sensitivity (CS normal=71.8%, CS50=89.1%, CS2F=94.6% and CS2.5F=97.12%). Notice that with increasing GR sensitivity, the 24 hr. cortisol AUC is lower than the normal cortisol profile, which is also in agreement with observations in PTSD subjects (Rohleder et al., 2004; Wahbeh and Oken, 2013; Wheler et al., 2006; Yehuda and Seckl, 2011). Moreover, the levels of IL6 and TNFα also increased consistently with increasing GR sensitivity suggesting a mechanistic association. The nonlinearity in the cortisol levels with respect to GR sensitivity is the function of the feedback inhibition parameters in the HPA immune axis, which may vary in populations leading to inconsistent data of cortisol levels.

**Figure 3:**
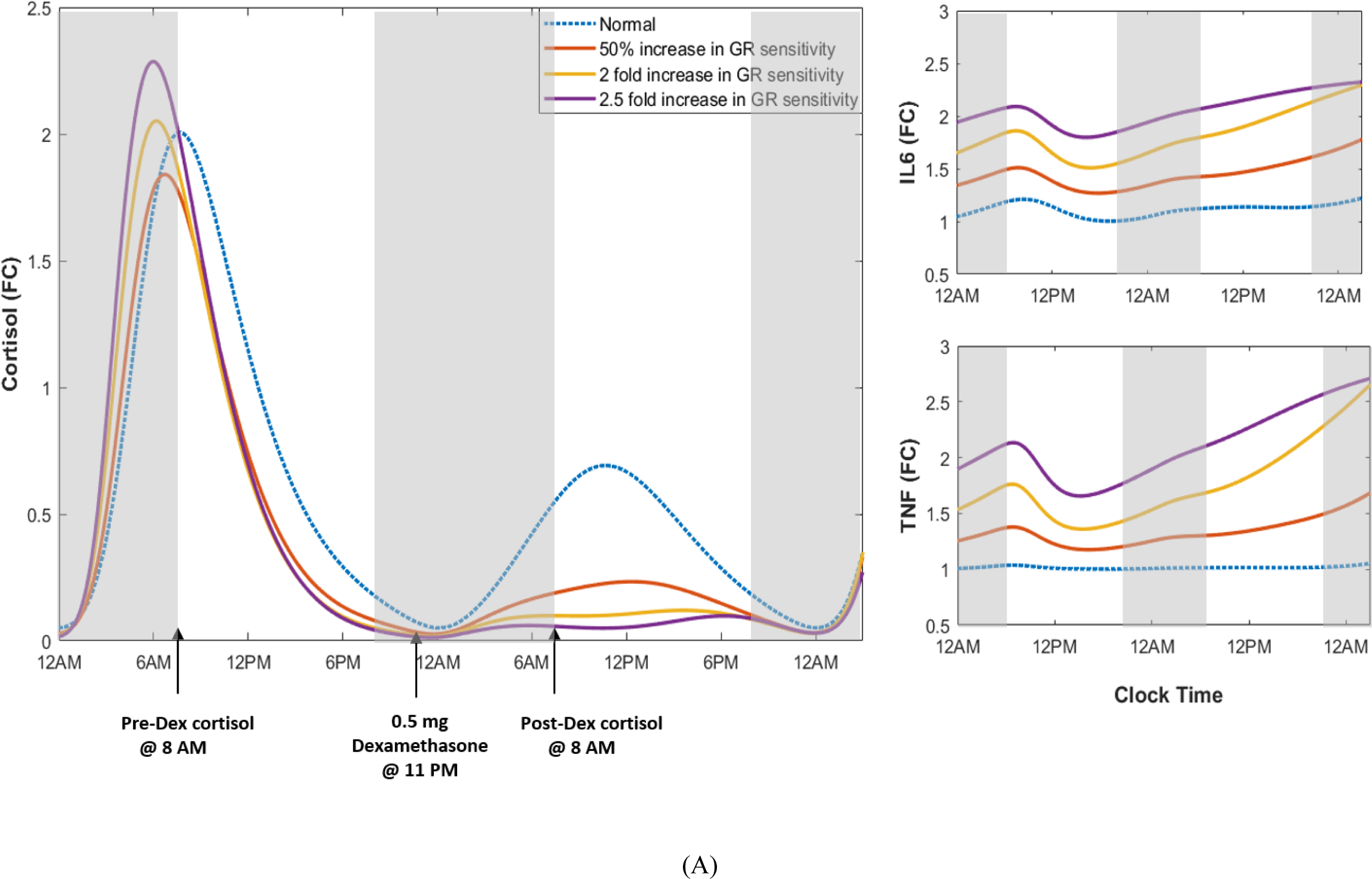
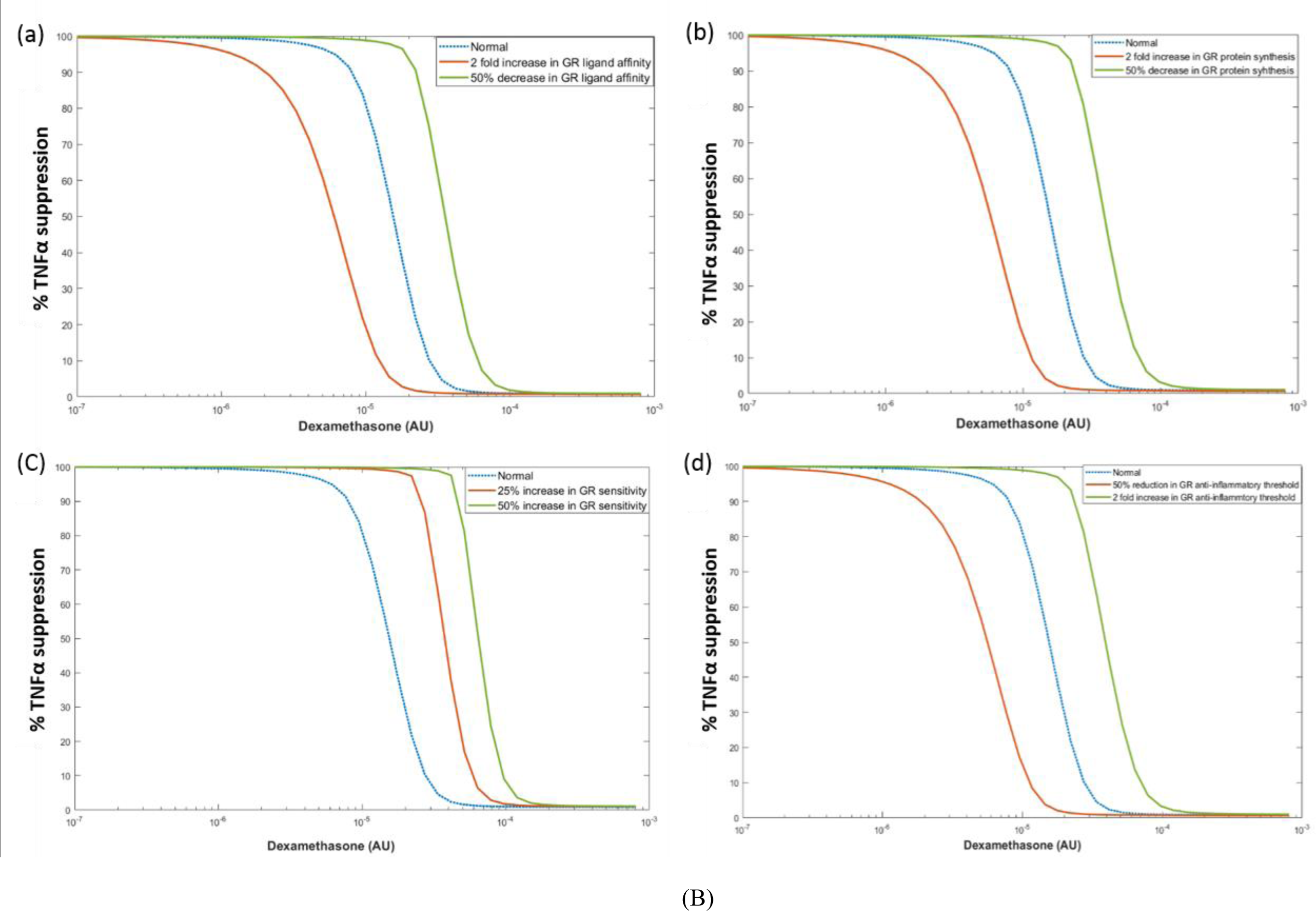
The simulated profiles of cortisol and cytokines for DEX suppression test for varying levels of GR sensitivity. The DEX dose was introduced at 11 PM. The cortisol values at 8 AM pre-DEX dose and at 8 AM post DEX dose were used to evaluate the percent of cortisol suppression in each case. It can be noted that the increase in cortisol suppression is consistent with increasing sensitivity irrespective of pre-DEX cortisol levels. Notice that increasing GR sensitivity is associated with consistent rise in cytokine levels alike cortisol suppression levels. The extent of cortisol suppression was 71.8%, 89.1%, 94.6% and 97.1% for normal, 50% increase, 2 fold increase and 2.5 fold increase in GR sensitivity, respectively. The peak cortisol levels decreased for up to 2 fold increase in GR sensitivity and increased on greater than 2 fold rise in GR sensitivity. The AUC for 24 Hrs. Cortisol was also reduced when GR sensitivity increased. (B) The simulated profiles for percent TNFα suppression by DEX dose. The model for inflammatory pathway was subject 25 ng/ml LPS stimulation and the TNFα suppression was recorded for varying doses of DEX (0 to 1000nM) incubated for 24 hrs. (a) The profiles for varying GR ligand affinity (*k_on* in Eqn.9 and 10). It can be noted that increasing GR ligand affinity decreases IC_50_-DEX level. (b) The profiles for varying GR protein synthesis (*ksynt_rp* in Eqn.9). It can be noted that IC_50_-DEX decreases with increasing GR synthesis. (c) The profiles for increasing GR sensitivity (Hill coefficients for GR feedback effects) in the inflammatory pathway. Increasing GR sensitivity increased the IC_50_-DEX levels. (d) The profiles for varying GR’s anti-inflammatory threshold (*q2* in Eqn.14)). Note that dotted curve represents the nominal curve with nominal anti-inflammatory threshold and sensitivity of the GRs, and the red and green curves are compared with the dotted-blue curve as the reference.

### Model simulations for IC_50_-DEX for cytokine expression

To analyze the sensitivity of the immune cells, we simulated the IC_50_-DEX test for TNFα expression using the mathematical model. The isolated sub-model for inflammatory pathway was stimulated with 25 ng/ml LPS and varying concentrations of dexamethasone concentrations (0 to 1000 nM) incubated for 24 hrs. The corresponding TNFα levels were recorded for different scenarios as shown in Figure 3B. Figures 3B (a) and 3B (b) shows the dexamethasone suppression of LPS-stimulated TNFα expression for 50% increase (red curve) and 50% reduction (green curve) in GR ligand affinity and GR protein synthesis, respectively; whereas the dotted blue curve represents the response for normal GR levels. Notice that the IC_50_ reduced with increasing GR ligand affinity (*k*_*on*_) and GR synthesis rate (*Vp*) indicating higher TNFα suppression as a function GR activity and abundance. This is consistent with the observations reported in PTSD subjects (Matić et al., 2013; Mirjam van Zuiden et al., 2011; Rachel Yehuda, 1991; Yehuda et al., 1995). Figure 3B(c) shows the dose-response curves for suppression of TNFα expression by varying levels of dexamethasone. The curves for 25% increase (red curve), normal (dotted blue curve) and 50% increase (green curve) in GR sensitivity (i.e. *nx* in Equation 14) are shown. We observed higher cytokine production on increasing GR sensitivity. It is noted that IC_50_ levels increased with increasing GR sensitivity, as observed in PTSD cases with increased IC_50_-DEX levels (de Kloet et al., 2007; Matić et al., 2013) along with an expected increase in the slope (steepness) of the curves. However, this is in contrast to our previous observations of reduced IC_50_-DEX lysozyme suppression and increased cortisol suppression in subjects with PTSD suggesting the prerequisite of increased GR availability or affinity for lowering IC_50_-Dex. Moreover, in the studies with increased IC_50_-Dex in PTSD subjects, there is a consistent finding of lower GR binding sites corroborating the relationship between GR abundance and IC_50_-Dex (de Kloet et al., 2007; Matić et al., 2013). We further varied the inhibitory threshold of GR on the inflammatory pathway (i.e. *q2* in Equation 14) (Figure 3B (d)). The figure shows IC_50_-Dex curves for a 2-fold increased (green) and 50% reduced (red) GR anti-inflammatory threshold, nominal (blue dotted). The curves indicate that IC_50_-Dex levels dictated the inhibitory thresholds.

### Differential changes in GR sensitivity of HPA-immune pathways

We simulated the effects separately of varying the parameters of the HPA-axis negative feedback and anti-inflammatory regulation of GRs. The model simulations reveal that increasing the sensitivity of only the negative feedback of HPA axis (*n,ny*) or reducing the inhibitory threshold (*Ki,Km1*) (without perturbing the anti-inflammatory GR effect) reduces the steady state levels of cortisol, ACTH and nuclear translocation of glucocorticoid receptors. The cytosolic GR increases for reducing inhibitory threshold and increasing sensitivity along with an increase in basal inflammation (See Figure 4: red curves for 50% increase in HPA-GR sensitivity, green curves-50% decrease in HPA-GR inhibitory threshold). Moreover, reducing sensitivity and increasing the inhibitory threshold, increased plasma levels of cortisol and ACTH, increased GR nuclear translocation and reduced the inflammatory response. However, reducing sensitivity reduced free GR levels and they were unaffected for increasing the inhibitory threshold (See Figure 4: yellow curve for 50% decrease in HPA-GR sensitivity, purple curve-50% decrease in HPA-GR inhibitory threshold). On the other hand, on increasing the sensitivity of inhibitory effect mediated by GRs on the inflammatory pathway (*nx*), there was increase in cortisol, ACTH and pro-inflammatory cytokines (TNFα and IL6) levels with decrease in free GR levels. However, decreasing *nx* or perturbing *q2* in either direction did not affect cortisol, ACTH or GR levels, but affected the basal cytokine levels (See supplementary Figure S1). Therefore, we hypothesize that changes in GR sensitivity of either pathway mechanistically influences inflammatory response.

**Figure 4.**
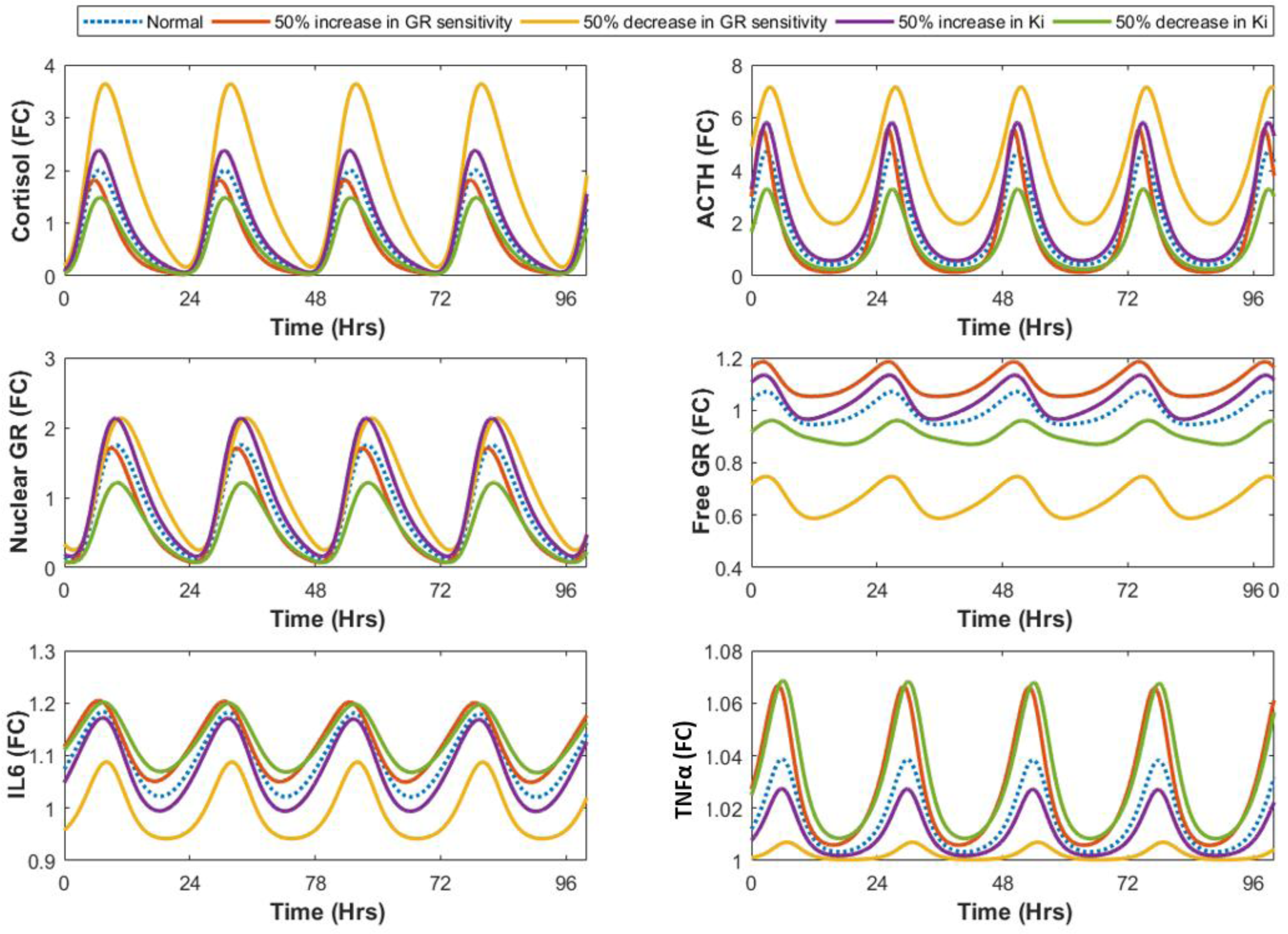
The simulated circadian profiles for varying the parameters for central negative feedback of GRs in the HPA axis. The Hill coefficient and the inhibitory thresholds for GR expression in Eqn. 1, 2, 8 and 9 were varied across the nominal values by 50%. It can be noted that increasing the strength of feedback regulation by decreasing inhibitory threshold or increasing the sensitivity decreases the ambient cortisol, ACTH and GR nuclear translocation which increases the basal TNFα and IL6 levels. On the other hand decreasing the sensitivity or increasing the inhibitory threshold increases cortisol, ACTH and GR translocation thereby suppressing basal inflammatory response.

### Causal Inference

To verify the mechanistic hypothesis of the increased GR sensitivity influencing the pro-inflammatory response, we performed causal association analysis of the relationship between the HPA axis measures with inflammatory cytokines. The covariate balancing propensity score (CBPS) weighting method was used to ensure the balance on the covariates. We also performed sensitivity analysis of the causal estimates (Christian Fong, 2018). The effects of HPA components were controlled for dexamethasone levels, age, BMI, education, race, ethnicity, smoking status, alcohol consumption, and medication use (7 categories: anti-depressants, sedatives, anti-convulsants, anti-allergy drugs, anti-inflammatories, anti-hypertensives, and pain medicines). The effects of NR3C1-1F promoter methylation were additionally controlled for unconverted/non-methylated cytosine levels. The un-adjusted Spearman correlation coefficients are reported in Table S4. The average causal effects and the corresponding robustness estimates between the two variables are displayed in Table S5. The average causal effects of the GR related variables on the cytokines and the corresponding robustness estimates (τ2) are shown in Figure 5A.

**Figure 5:**
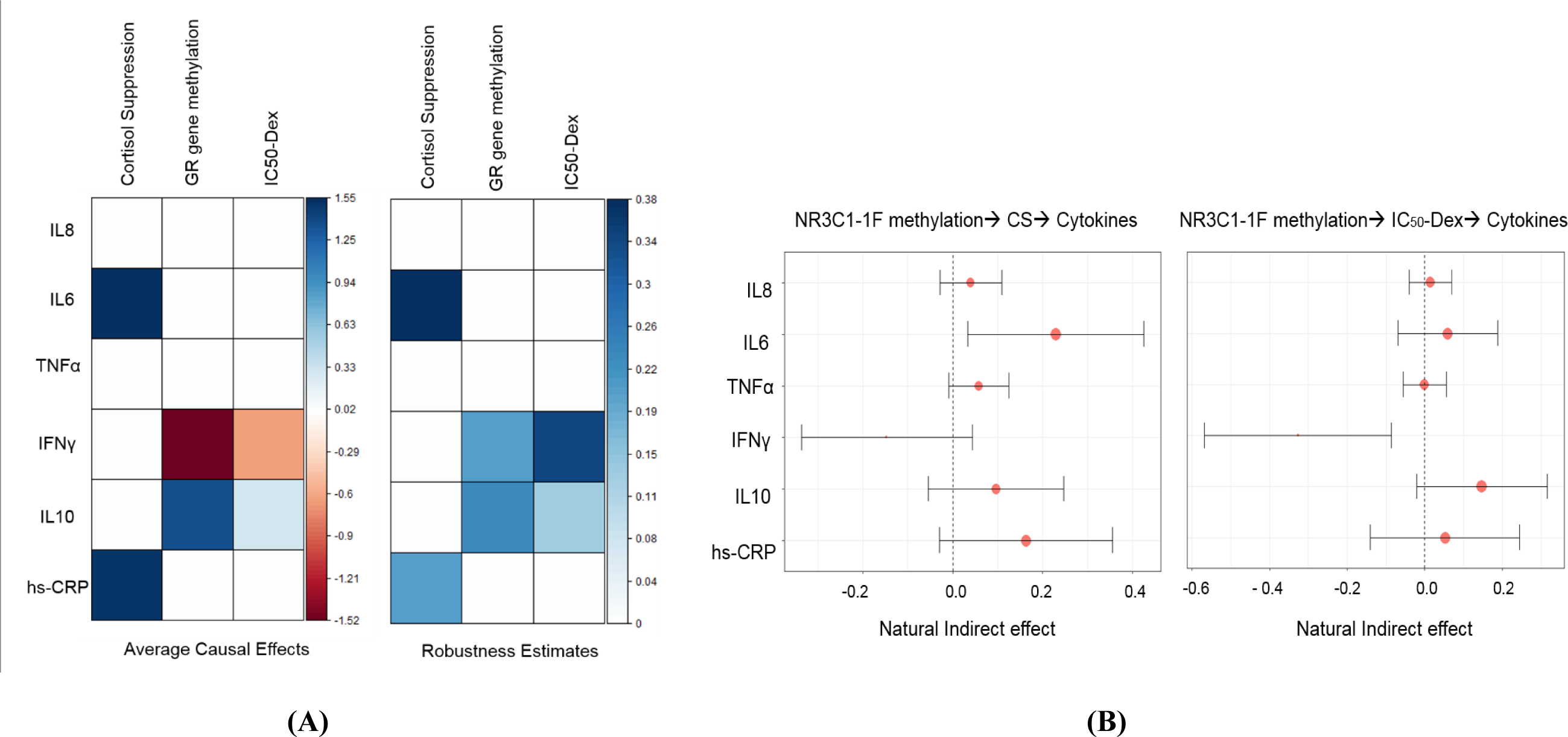
(A) Average causal effects and corresponding robustness estimates for cortisol suppression, GR gene promoter methylation and IC_50_-DEX on cytokines. It is noted that cortisol suppression is significantly causally associated with IL6 and hs-CRP expression. NR3C1-1F promoter methylation and IC_50_-DEX for lysozyme suppression are negatively associated with IFNγ and positively associated with IL10. All the causal associations are robust and less sensitive to unknown confounding. Cortisol suppression and NR3C1-1F promoter methylation showed a trend level effect on TNFα (See Table S5). (B) Forrest plots for the causal mediated effects of GR gene promoter methylation on cytokines through cortisol suppression and IC-_50_ Dex for lysozyme suppression evaluated by the natural effects models. The effect of NR3C1-1F promoter methylation on IL6 is statistically significant through cortisol suppression; and the effect on IFNγ is significant through IC_50_-DEX. These suggests that the changes in GR sensitivity may be due to lower NR3C1-1F promoter methylation that would increase GR expression leading to lower IC_50_-Dex levels and increased cortisol suppression, which further is associated with pro-inflammatory response.

Since we observed significant correlations and causal association of NR3C1-1F promoter methylation with cortisol decline post dexamethasone dose, IC_50_ and inflammatory cytokine levels, we further tested the causal mediation hypothesis on the effect of GR methylation on pro-inflammatory cytokines mediated by changes in GR sensitivity, using a natural effects model (VanderWeele, 2016). The results of the mediation analysis are reported in Table S6. A trend level mediated effect was noted on IL6, TNFα and hs-CRP. A significant causal mediation effect of GR through IC_50_-DEX was also noted for IFNγ and a trend in effect on IL10. These results indicate a trend in the effect of changes in NR3C1-F promoter methylation on GR sensitivity and subsequent inflammatory response. The causal mediated effects and the corresponding confidence intervals measured by natural indirect effects are shown in Figure 5B.

## Discussion

### Mechanistic insight on GR sensitivity and inflammation

To further analyze the results from the simulations, we plotted the feedback gains of GR and cytokine mediated effects in the HPA-immune axis (Figure 6A), namely, the anti-inflammatory activity of GR on the inflammatory pathway, the negative feedback activity of GR on the HPA axis and its own synthesis, and the activation effects of pro-inflammatory cytokines on the HPA axis (See Figure 6B). Notice that increasing GR sensitivity decreases the anti-inflammatory effect (Figure 6B (a)), increases the activation of HPA axis by TNFα (Figure 6B (b)), increases the inhibitory effect of negative feedback (Figure 6B (c)) and decreases negative and positive feedback on its own synthesis (Figure 6B (d)). The competing effect of these multiple feedback mechanisms would determine the resultant states of the variables in the HPA-immune network.

**Figure 6.**
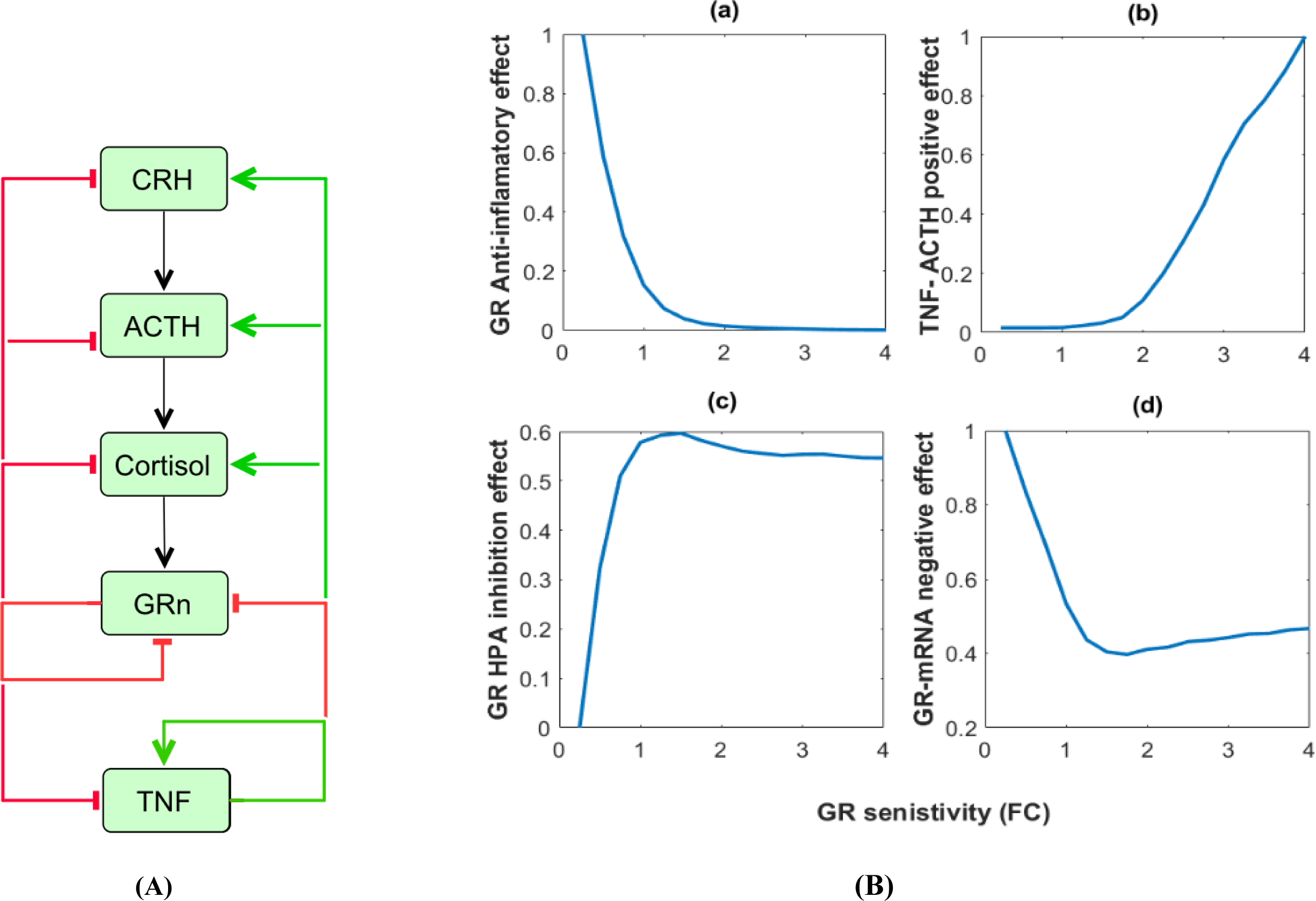
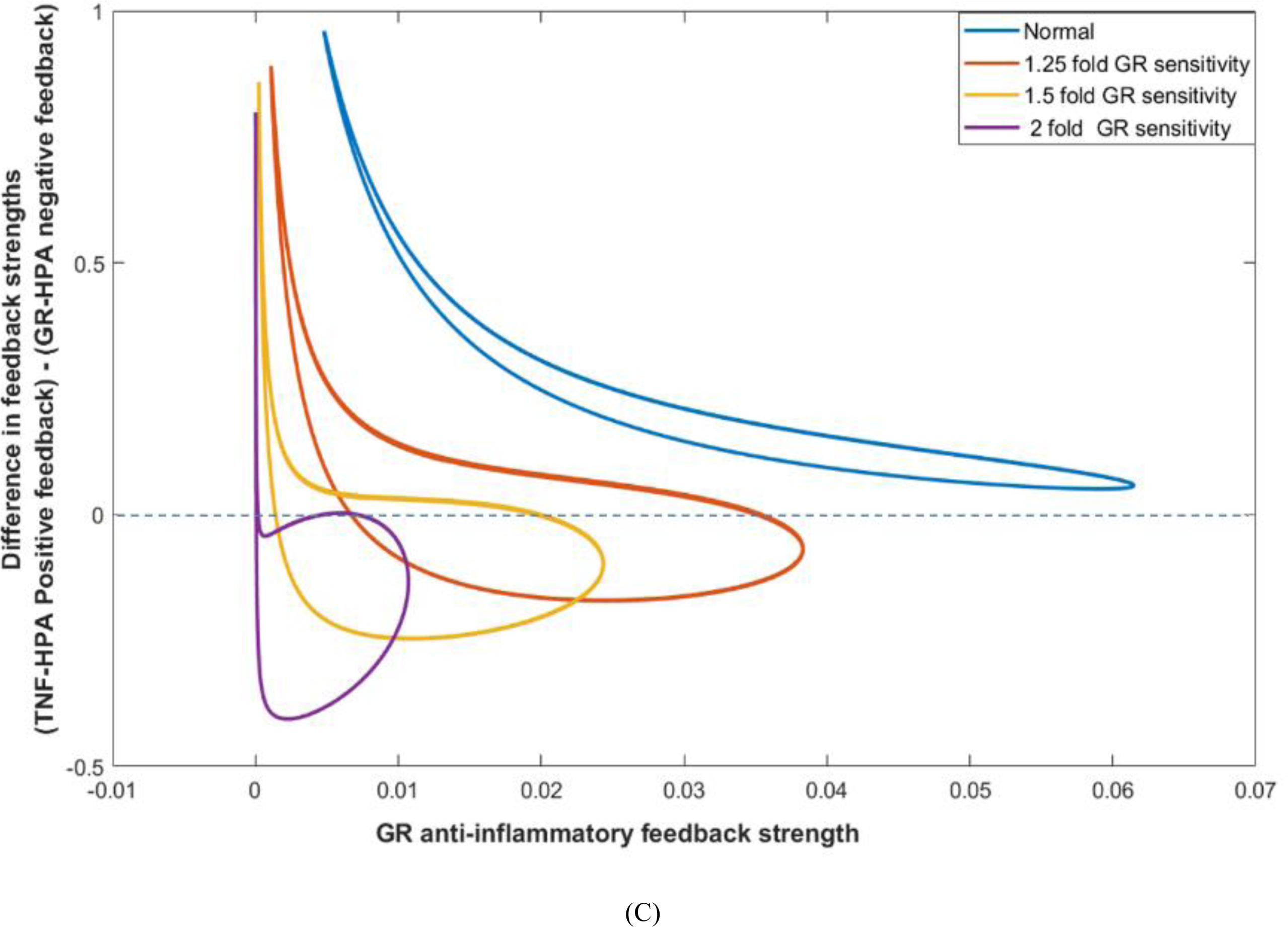
(A) The feedback regulatory system in the HPA-immune axis, wherein GR inhibits its own synthesis and downregulates cytokines (TNFα), whereas, TNFα activates CRH, ACTH and cortisol release. (B) The glucocorticoid receptors and cytokine related feedback gains for varying GR sensitivity. The values are scaled with their respective maximum values. (a) Anti-inflammatory activity of GR (Eqn. 14); (b) Activation of ACTH secretion by TNFα (Eqn. 2); (c) Negative feedback of GRs on the HPA-axis; (d) Negative feedback of GR on its mRNA synthesis (Eqn.8); (C) The phase-plane representation of the circadian dynamics of the feedbacks in the network motif. The difference in the strengths of TNFα mediated positive feedback (expression in green in Eqn. 2) and GR-mediated central negative feedback (expression in red in Eqn.2) are plotted with respect to GR-mediated anti-inflammatory feedback. The limit cycles are plotted for different levels of increasing GR sensitivity coefficients. It should be noted that with increasing sensitivity coefficients the extent of GR anti-inflammatory effect decreases, whereas the net difference between the strength of TNFα-mediated positive effect on the HPA axis and the GR negative feedback on HPA axis becomes negative, indicating inhibition of cortisol production due to the dominance of GR-negative feedback on the HPA axis. However, the relative amplitude of TNFα positive feedback effect increases with increasing sensitivity.

A simple interpretation might suggest a decrease in the inflammatory response with higher GR sensitivity, however, there are two reasons why this could not be the case; first is the effect through the central negative feedback that reduces the gain on GR nuclear translocation due to lower ACTH stimulated cortisol release(ligand required for GR binding) and second is the subsequent reduction in the anti-inflammatory response due to realization of increase in the inhibitory saturation threshold because of the higher sensitivity parameter that appears as the power of inhibitory threshold (*q*2*^nx^*) in the saturation kinetics (See Equation 14) (because *q2*≫*GRn*). Therefore, the activated inflammatory response acts to further upregulate the HPA axis in an attempt to restore homeostasis, however at the cost of a dysregulated immune response with increased pro-inflammatory milieu. This can be viewed as an adaptive response to normalize inflammation. However, it appears that the cytokine-induced cortisol release would not be sufficient to achieve perfect adaptation due to relative differences in the activation threshold of cytokines (*q5 in Eqn.2*) and inhibitory threshold of GR (*Ki in Eqn.2*) on HPA-axis, wherein (*q5*≫*TNFα basal*) and (*Ki*≪*GRn basal*) suggesting a stronger negative feedback by GRs as compared to the positive effect of cytokines. Therefore, a relatively lower cortisol would lead to a relatively lower GR anti-inflammatory activity as compared to that would have been required for a corresponding rise in cytokine levels. This can be noticed from the Figure 6C, wherein the limit cycles for relative difference in TNFα-positive feedback and GR negative feedback on HPA-axis becomes negative, indicating dominance of GR-HPA negative feedback over TNFα-HPA stimulatory effect and a corresponding decrease in the amplitude of GR-anti-inflammatory effect, with increase in GR sensitivity. Moreover, increased TNFα downregulates cytokine GR activity, inducing a positive loop for pro-inflammatory state (See TNFα feedback in Eqn. 10 and Eqn. 11) (Webster et al., 2001). These mechanisms can be supported from our data with statistically significant causal association between pre-DEX cortisol levels with the pro-inflammatory cytokine IFNγ (ϒ=0.747, p=0.030, q=0.057, τ_2_=0.118); and a negative association with anti-inflammatory cytokine IL10 (ϒ=-0.712, p=9.4E-3, q=0.002, τ_2_=0.201) and IC_50_-DEX (ϒ=-0.386, p=0.017, q=0.039, τ_2_=0.196).

The inhibitory effects of GC on inflammation are mainly observed for the pharmacological doses of GC, whereas at physiological GC levels, the effect are subtle. This implies that the inhibitory threshold (*q2*) of the dose-response curve is elevated above the ambient GC levels; therefore increasing the sensitivity of GRs *(nx)* will decrease the overall rate of inhibition for the ambient levels of GC (Eqn.14) (*GRn<q2*); however it sensitizes the effect on additional GC exposure with a steep response on realizing the saturation threshold when *GRn=q2* (See Figure S2). Corroborating these mechanisms, we observed causal association of post-DEX cortisol levels with IL6 (ϒ= −0.199, p=1.1E-4, q=2.3E-4, τ_2_=0.3775) and a negative trend with hs-CRP; and an association between extent of decline in cortisol levels with hs-CRP (ϒ=0.697, p=7E-3, q=0.016, τ_2_=0.193) and a negative trend with IL10. Moreover, cortisol suppression was positively causally associated with IL6 (ϒ=1.55, p=3.0E-4, q=5E-4, τ_2_=0.375) and hs-CRP (ϒ=1.5, p=0.019, q=0.03, τ_2_=0.205).

### Implications of IC_50_-DEX and GR promoter methylation to inflammation in PTSD

The analysis of simulated IC_50_-DEX curves reveals that IC_50_ levels are indicative of the amplification in the anti-inflammatory effects of GC, whereas the slope of these curves represents the sensitivity of the anti-inflammatory effect. Therefore, the slopes of the IC_50_-dex curves should correlate with cortisol suppression for systemic changes in GR sensitivity. However, IC_50_ levels may or may not coincide with the GR sensitivity of the negative feedback of the HPA axis. This can also be noticed from the variability of correlation between cortisol suppression test and IC_50_-lysozyme suppression test in different PTSD samples (Matić et al., 2013; Rohleder et al., 2004; Yehuda et al., 2003). IC_50_-DEX is also indicative of GR receptor number or Bmax levels as observed in other reports on PTSD subjects (de Kloet et al., 2007; Matić et al., 2013), which is consistent with the observation that increased GR density and dimerization potential are responsible for higher GR activity (Robertson et al., 2013).

Supporting these observations in our data, we noted a statistically significant positive causal association between NR3C1-1F promoter methylation and IC_50_-DEX assay (ϒ=0.877, p=4.7E-3, q=0.01, τ_2_=0.242) in our sample, suggesting lower methylation (indicative of increased GR expression) may be responsible for a lower anti-inflammatory GR threshold. Consistent with the model simulations of increased pro-inflammatory cytokines and reduced IL10 (anti-inflammatory cytokine) with increasing GR sensitivity, IC_50_-DEX in our data for lysozyme suppression showed a negative causal association with IFNγ (ϒ= −0.646, p=3.34E-4, q=1.8E-3, τ_2_=0.338) and a positive association with IL10 (ϒ=0.29526, p=0.043, q=0.23, τ_2_=0.132). Moreover, we noticed identical effects of GR methylation on the cytokines IFNγ (ϒ= −1.516, p=4.7E-3, q=0.01, τ_2_=0.204) and IL10 (ϒ=1.33, p=3.25E-3, q=8.6E-3, τ_2_=0.238) corroborating the hypothesis of increased GR availability or binding affinity could be responsible for lower IC_50_-Dex and increased inflammation. These analyses also supports the hypothesis on sensitization of pro-inflammatory response due to increased GC receptor occupancy inferred from studies on temporal dose-response for cortisol and inflammatory response in animal models (Frank et al., 2010).

Furthermore, the methylation of NR3C1-1F promoter showed a negative causal association with measures of the HPA axis: pre-DEX cortisol levels (ϒ=-0.596, p=8.5E-6, q=5.22E-5, τ_2_=0.371), cortisol difference (ϒ=-0.80, p=9.78E-6, q=5.22E-5, τ_2_=0.379), pre-DEX ACTH levels (ϒ=-0.455, p=8.25E-4, q=3.3E-3, τ_2_=0.286), ACTH difference (ϒ=-0.734, p=0.011, q=0.022, τ_2_=0189) indicating higher GR activity (availability) with reduction in methylation (Yehuda et al., 2015). The trend level significance in the causal mediation analysis (See Table S6) suggests that the lower methylation of NR3C1-1F promoter may affect GR sensitivity and contribute to a pro-inflammatory response in PTSD.

Although, lower IC_50_-DEXsuggests a higher anti-inflammatory response; it must be noted that the IC_50_-DEX levels are in-vitro characterization evaluated in the PBMCs and may or may not be representative of the systems level response of HPA-immune network due to disconnection between HPA axis regulatory mechanisms. In the in-vivo scenario, the increased GR sensitivity would increase GR expression levels (See Figure 4-Free GR), as well as pro-inflammatory cytokines induce GR expression (Webster et al., 2001), which in effect could yield lower IC_50_-DEX lysozyme suppression in the PBMCs evaluated in-vitro (See Figure 3B (b)). However, this may not directly translate to the increased in-vivo anti-inflammatory potential of GRs.

### Plausible mechanism for variability of HPA features in PTSD cohorts

In a typical biological system, the sum of the different regulatory mechanisms is determined by the parameters of saturation kinetics, and the homeostatic steady state level is determined by the relative values of saturation thresholds with respect to the steady sate values of activator or inhibitor concentrations. Therefore, changing a steady state due to a perturbation in one feedback element can change the relative gain from other feedback components due to shifts in the regulatory landscape with respect to the saturation thresholds and respective sensitivity parameters. In view of such dynamics, resistance or sensitivity in one feedback can affect the gain and response of other feedback loops thereby shifting the steady state of the system away from homeostasis. Such a scenario is likely in the case of the HPA-immune axis and may underlie the variation in the findings on HPA state variables in different populations.

To analyze such a scenario, we simulated the model for varying levels of GR sensitivity and the inhibitory threshold in the feedback loop enacted by GRs (represented by expressions in Equations 1, 2, 8 and 14). To simulate the systemic GR effects, the simultaneous variations in both the central (*n, ny* and *Ki, Km1*) and immune (*nx* and *q2*) negative feedback regulation were considered for this analysis. The corresponding states for cortisol, ACTH, nuclear GR, cytosolic GR, IL6 and TNFα are plotted in Figure 7A. It is noted that the cortisol, ACTH and GR nuclear translocation levels increase with increasing inhibitory saturation thresholds and decrease nonlinearly with increasing GR sensitivity, whereas cytosolic GR levels increase nonlinearly with increasing sensitivity and inhibitory threshold. However, the inflammatory cytokine levels increase with an increase on either axis. To visualize effect of varying strengths of GR negative feedback and the excitatory effect of cytokine on HPA-axis, we plotted the HPA-immune axis variables, varying the feedback thresholds of GR negative feedback (*Ki, Km1* and *q2*) and the cytokine-HPA axis activation effect (*q5*) (Figure 7B). It was observed that the steady-state levels of all features of HPA-immune axis increased with increasing the inhibitory threshold due to decreased feedback inhibition. Whereas, decreasing the activation threshold of the HPA axis by TNFα resulted in increased levels of cortisol, ACTH and GR nuclear translocation and decreased the basal levels of cytokines and free GR levels.

**Figure 7.**
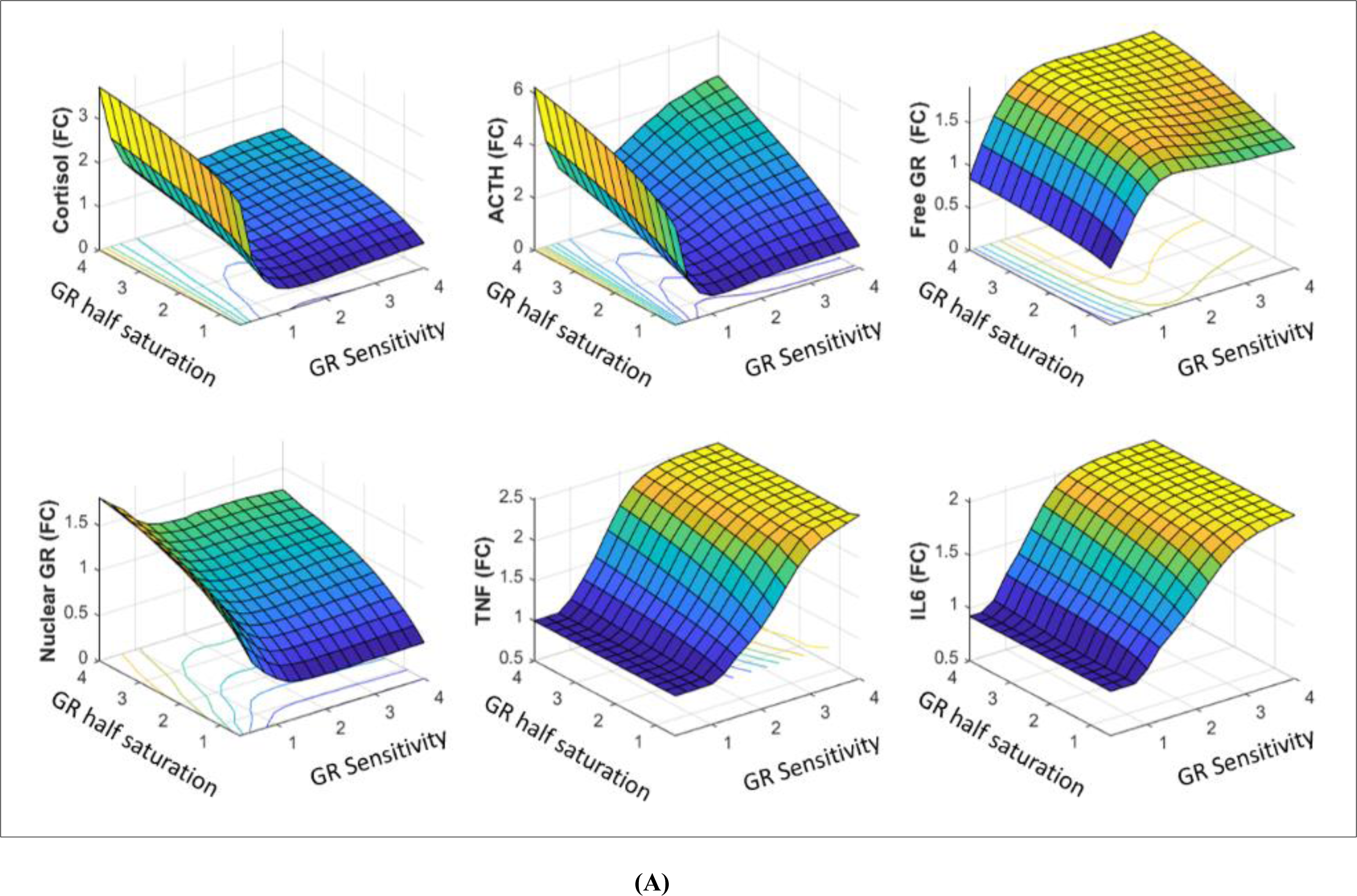
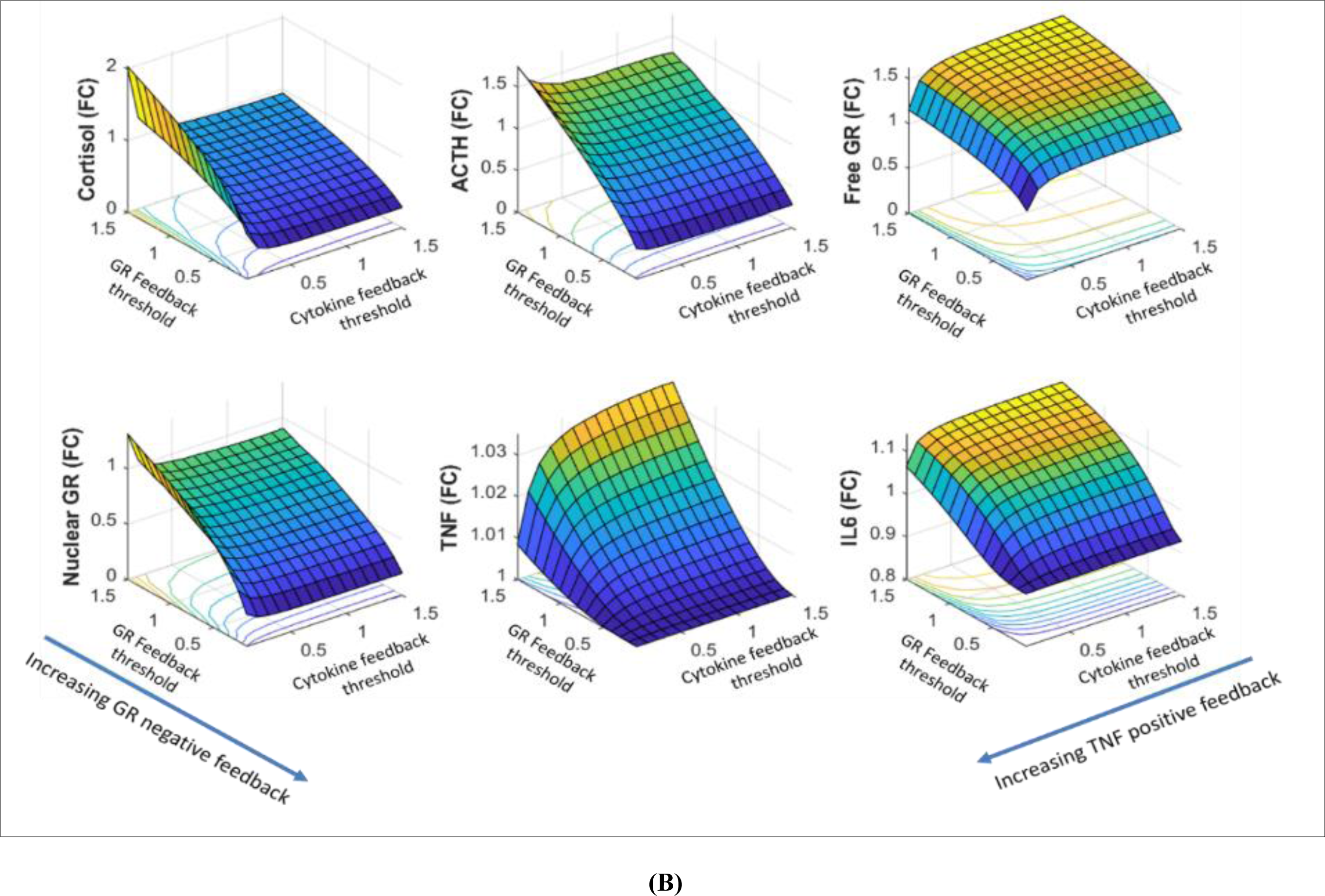
(A) 3D plots of the system steady states obtained through model simulations by varying the inhibitory threshold and the sensitivity parameters of the feedback loops in HPA-immune axis (overall feedback inhibition is shown in red blunted edges in Fig. 6A). It is noted that GR nuclear translocation decreases with increasing sensitivity thereby increasing free cytosolic GR. GR nuclear translocation follows the profiles for cortisol wherein it increases with increasing the saturation threshold. Pro-inflammatory response increases on increasing both inhibitory saturation threshold and sensitivity. It can be noted that all the states are elevated with increasing both sensitivity and inhibitory threshold except an opposite trend in cytosolic GR. (B) 3D plots of the system steady states for varying the feedback thresholds for activation of the HPA axis by cytokines (*q5* in Eq.14) and inhibition of HPA-immune axis by GRs (*Ki* in Eq. 1, 2 and 8). Increasing the inhibitory threshold of GR negative feedback increases the values of the variables in the HPA axis and the basal levels of cytokines, whereas decreasing the activation threshold of cytokines for HPA axis increases cortisol, ACTH and GR translocation and decreases the free GR and basal cytokine (TNFα and IL6) levels.

At the physiological level, while exposure to traumatic experience is associated with an acute increase in cortisol, the negative feedback dampens the release of cortisol to maintain homeostasis. However, if an increased cortisol production driven by inflammatory cytokines modulating the HPA axis is sustained due to varied reasons other than reduced HPA axis negative feedback, further expression of GR could be induced rendering a shift in the basal inhibitory threshold. Such a state can be viewed as glucocorticoid resistant state to inflammation with increased glucocorticoid levels (Cohen et al., 2012). In the Figure 7A, it can be observed that the surface of the GR nuclear translocation is nonlinear with respect to GR sensitivity, wherein reducing sensitivity increases GR translocation due to increased ligand availability, and increasing the sensitivity would increase GR translocation because of increased inflammatory response. However, on further increasing the sensitivity, the GR translocation subsides due to lack of glucocorticoid synthesis and ligand availability. Due to such a nonlinear effect of GR sensitivity on the activity of the HPA-immune axis, it can be expected that one would observe varied effects of HPA dysregulation (de Kloet et al., 2007; Gola et al., 2014) in different cohorts of individuals with PTSD.

Some set of predefined factors such as genetic predisposition, childhood trauma and inflammatory susceptibility might determine the basal parameters of the HPA axis and immune regulation in respective individuals (Lian et al., 2014), which in turn determine its robustness. The variability in the cortisol levels observed in other studies can be possibly accounted by such an individual level differences in parameter space and associated inflammatory state. The analysis suggests an increased levels of inflammation, cortisol suppression and plasma cortisol (as observed in our data) can be achieved for a shift in parameter space in the diagonal direction on the 3D plot (Figure 7A) (i.e., increase in both sensitivity parameter and the inhibitory threshold of negative feedback) resulting in a reduced GR negative feedback strength at ambient GR levels..

## Conclusion

Our analysis suggests that the increased GR sensitivity is mechanistically associated with an increased inflammatory response in individuals with PTSD. Increased GR sensitivity would lower the contribution of ACTH stimulated cortisol release due to higher GR negative feedback effect, which would increase pro-inflammatory response and subsequent cytokine induced cortisol secretion to normalize inflammation. However, due to dominance of GR mediated negative feedback over cytokine mediated positive feedback on the HPA-axis, the cortisol rise is insufficient to overcome regulation by precursors of an inflammatory response. On the other hand, the downregulation of GR activity by pro-inflammatory cytokines would reduce its anti-inflammatory effect at basal GR levels. Further, on selectively increasing the GR sensitivity of anti-inflammatory effect, the dose response curve turns sigmoidal; wherein, although the GR anti-inflammatory effect is higher at higher GR levels, the GR activity is reduced at lower GR concentrations as compared to the normal GR dose-response due to higher anti-inflammatory threshold of GRs relative to its ambient levels. This may lead to lower anti-inflammatory effect and an increased pro-inflammatory response (See Figure S2, TNFα and IL6) despite normal GR levels.

The results of the IC_50_-DEX lysozyme suppression test characterize an amplification in the anti-inflammatory effect of glucocorticoids and could be indicative of GR ligand binding affinity or GR expression levels. We also noticed a trend in causal mediation effects between lower methylation of NR3C1-1F promoter regions on inflammatory cytokines through increased GR sensitivity assessed by IC_50_-Dex and post-DEX cortisol levels, suggesting that the methylation of NR3C1-F promoter may affect GR sensitivity and its anti-inflammatory potential. However, the specific molecular mechanisms that contribute to increased GR sensitivity and reduced IC_50_-Dex remains elusive and need further research. Our analysis motivates future investigations on assessing the role of GRs in controlling inflammatory response.

## Author contributions

PRS conceived the work, performed the analysis and wrote the manuscript. SM and OW generated the cytokine data and supervised the manuscript. FJ and RY generated the neuroendocrine data and supervised the manuscript. DA and CM recruited the study.

## Acknowledgement

This work was supported by a research grant from U.S Army Medical Research and Material Command (USAMRMC) under award number W911NF-17-2-0086. We are thankful to Dr. Owen Wolkowitz and Dr. Michael Meaney for their comments and edits on the manuscript. We also acknowledge Dr. Matthew Kobe and Dr. David Jackson from US Army for editing and reviewing the manuscript.

## Disclaimer

The views, opinions, and findings contained in this report are those of the authors and should not be construed as official Department of the Army position, policy, or decision, unless so designated by other official documentation. Citations of commercial organizations or trade names in this report do not constitute an official Department of the Army endorsement or approval of the products or services of these organizations.

## Materials and Methods

### Participants

The sample and procedures have been described previously and in full detail (Lindqvist et al., 2014; Yehuda et al., 2015). Briefly, the data described in this report were derived from a sample of 165 US male veterans who were deployed to Iraq or Afghanistan (81 without PTSD, 81 with PTSD). Participants were recruited from the James J. Peters Veterans Affairs Medical Center (JJPVAMC)/Icahn School of Medicine at Mount Sinai (ISMMS), and New York University (NYU) Langone Medical Center (NYULMC)/NYU School of Medicine (NYUSM) through advertising in the clinic (VAMC) and community (NYU). The study was approved by the IRBs of the JJPVAMC, ISMMS, NYULMC and NYUSM; all participants provided written, informed consent.

### Clinical assessment

Exclusion criteria included a recent history of alcohol dependence or drug abuse/dependence; lifetime history of a psychotic disorder, bipolar disorder, or obsessive-compulsive disorder; current exposure to recurrent trauma; prominent suicidal or homicidal ideation; neurological disorder or systemic illness affecting central nervous system function; and recent initiation (< 2 months) of psychiatric medication, anticonvulsants, antihypertensive medication, or sympathomimetic medication. Psychiatric diagnostic information based on DSM-IV criteria, including PTSD diagnosis, was determined by doctoral level psychologists using the Structured Clinical Interview for DSM-IV (SCID) and the Clinician Administered PTSD Scale (CAPS). All participants reported exposure to a combat-related DSM-IV PTSD Criterion A trauma. Veterans who did not meet lifetime diagnostic criteria for PTSD and had a CAPS score ≤ 20 were included in the PTSD-group. Veterans who met DSM-IV diagnostic criteria for PTSD and had a CAPS score ≥ 40 were included in the PTSD+ group. Combat veterans with CAPS scores between 20 and 40 were not included in the study. Several self-reported clinical assessments were also administered to the sample including the PTSD Checklist for DSM-IV (PCL), the Mississippi Combat Scale (MCS) the Pittsburgh Sleep Quality Index (PSQI) and the Early Trauma Inventory (ETI).

### Neuro-endocrine Assays

#### Cortisol

levels in plasma were assayed by Cortisol ELISA Kit from IBL-America (Minneapolis, MN). It is a solid phase enzyme-linked immunosorbent assay, based on the principle of competitive binding. The microtiter wells were coated with a monoclonal antibody directed towards an antigenic site on the cortisol molecule. Endogenous cortisol from an unknown competes with a cortisol-horseradish peroxidase conjugate for binding to the coated antibody. After incubation the unbound conjugate was washed off. The amount of bound peroxidase conjugate is inversely proportional to the concentration of cortisol in the unknown. After addition of the substrate solution, the intensity of color developed is inversely proportional to the concentration of cortisol in the unknown. Assay sensitivity: 2.5 ng/mL. The intra-assay and inter-assay coefficients of variation for this assay 5.3% and 9.8%, respectively. Two blood samples were assayed for the determination of cortisol before and after DEX administration. Decline of cortisol from Day 1 to Day 2 was used as a measure of DEX suppression.

#### ACTH

levels in plasma were assayed by using ACTH ELISA kit (ALPCO Diagnostics, Windham NH). In this assay, calibrators, controls, or patient samples were simultaneously incubated with the enzyme labeled antibody and a biotin coupled antibody in a streptavidin-coated micro plate well. At the end of the assay incubation, the microwell was washed to remove unbound components and the enzyme bound to the solid phase was incubated with the substrate, tetramethylbenzidine (TMB). An acidic stop solution was added to stop the reaction and convert the color to yellow. The intensity of the yellow color is directly proportional to the concentration of ACTH in the sample. A dose response curve of absorbance unit vs. concentration was generated using results obtained from the calibrators. Concentrations of ACTH was determined directly from this curve. Assay sensitivity: 0.5 pg/mL. The intra-assay and inter-assay coefficients of variation for this assay 5.7% and 8.0%, respectively. Two blood samples were assayed for the determination of ACTH before and after DEX administration. Decline of ACTH from Day 1 to Day 2 was used as a measure of DEX suppression.

#### Lysozyme Activity Assay

The test for examining the inhibition of lysozyme synthesis and release was carried out in 96-well culture plate in a total volume of .22 mL, modified from Panarelli et al (1994).Lysozyme activity was measured by turbidimetric method using *Micrococcus lysodeikticus* (Sigma) as the substrate. *Micrococcus lysodeikticus* was prepared in 0.1 mol/L phosphate buffer, pH 6.3, at a concentration of .05% and homogenized with a tissue grinder equipped with a Teflon pestle (Wheaton, St. Millville, New Jersey) with 3 strokes. 20 µL of supernatant of cell culture was incubated with 150 µL of substrate in a 96-well plate at 37°C for 7–10 min with shaking and then kinetically read by a microplate reader at 450 nm for 20 min. Cells (3.5-4.0 × 105) were incubated with 0, .5, 1, 2.5, 5, 10, 50, 100, and 200 nmol/L of DEX (Sigma) at 37°C in a humidified atmosphere with 5% CO2 for 3 days. Each concentration of DEX was incubated in triplicate. After centrifuging the plate, 120 µL of supernatant were removed and pooled from each triplicate well. The standards were prepared using pure lysozyme from chicken egg white (Sigma) dissolved in RPMI-1640 as used for the cell culture. The inhibition curve was drawn as concentration of DEX versus relative activity of lysozyme. Results were expressed as IC_50_-DEX (nmol/L) based on the concentration of DEX at which 50% of lysozyme activity was inhibited. The intra- and inter-assay coefficients of variation for the measurement of lysozyme activity were 6.9% and 9.8% respectively (Yehuda et al., 2004).

#### DNA cytosine methylation of the NR3C1-1F promoter

In order to quantify this epigenetic marker, genomic DNA was extracted from the frozen PBMC pellets following the Flexigene DNA kit protocol (Qiagen, CA, USA). Methylation mapping of the 39 CpG sites in the NR3C1-1F promoter was performed as previously described by our collaborator Dr. Micheal Meaney at McGill University using 30 clones per sample. Briefly, sodium bisulfite conversion was carried out according to the EpiTect Bisulfite kit protocol (Qiagen) using 0.8 µg of genomic DNA in each conversion reaction and 0.8 µg of Universal Methylated Standard (Zymo Research, CA, USA) to check completion of the reaction. The genomic region of the human GR exon 1F promoter was subjected to PCR amplification using the following primer sequences: 5’-GTG GTG GGGGAT TTG-3’ (forward); 5’-ACCTAATCTCTCTAAAAC-3’ (reverse) following previously published procedures (3, 16). The resulting PCR product was subjected to another round of PCR, using the following nested primers: 5’-TTTTTGAAGTTTTTTTAGAGGG-3’ (forward); 5’-AATTTCTCCAATTTCTTTTCTC-3’ (reverse) following previously published procedures. The resulting PCR products were analyzed on a 2% gel agarose gel and then purified using QIAquick kit (Qiagen). The PCR products were subcloned using a PCR product cloning kit (Qiagen) and individual plasmid containing the ligated promoter regions were extracted and sequenced (Genewiz, NJ, USA). The sequences for 30 individual clones were aligned and analyzed in the DNA Alignment software program BioEdit (Ibis Biosciences, USA). The DNA samples were analyzed in batches of 20 to 30 samples. Variability in the DNA bisulfite treatment did not exceed 2% between the batches. Two measures were calculated: first, the number of clones with at least one methylated CpG site divided by the total number of clones was calculated to yield an estimate of the percentage of methylated clones; second, the number of methylated clones (out of 30) at each of the 39 CpG sites was converted to a percentage, and percentages across sites were summed to create a total percentage of methylation across the NR3C1-1F promoter sequence.

#### Cytokine Assays

The blood samples were drawn in the morning after overnight fasting. The serum levels of cytokines (IL6, TNFα, IFNγ, IL10 and high sensitive C-reactive protein (hs-CRP) were estimated as per the methods reported in our earlier reports (Lindqvist et al., 2014).

#### Statistical analysis and correlations

To determine the group differences in the features of the HPA-axis and inflammatory pathway, we performed a Wilcox test using the raw data. The p-value threshold of α=0.05 was considered as statistically significant. The q-values for the correlations and causal analysis were evaluated using q-value package in R (Storey et al., 2017). We performed correlational analyses to determine correlations between the features of the HPA axis and cytokines using spearman correlation coefficients. The package Hmisc was used for correlational analysis in R.

### Average causal effects

For the causal analysis, the clinical data was imputed for missing values, log transformed and median normalized. We used covariance balancing propensity scores for weighting in the regression analysis for estimation of average causal effects (ACE). The CBPS package (Christian Fong, 2018) in R was used for causal analysis. A non-parametric estimation of covariance balancing propensity scores was performed to obtain weights for evaluating average causal effects. The population average causal effects were estimated using combined data for controls and cases with adjustments for group effects. We also performed sensitivity analysis to estimate the confidence on the causal estimates using the treatSens package (Carnegie et al., 2016) in R. The sensitivity analysis estimates the extent of bias required to nullify the causal effect rendering it to be insignificant. The standardized bi-parametric sensitivity estimate curves were obtained. The intersection point of the X=Y line with the sensitivity curves were evaluated as the measure of robustness of an association. The intersection of the X=Y line with the curve where ACE=0 represented by τ_1._ The intersection of the X=Y line with the curve where ACE is insignificant (p>0.05) is represented by τ_2._ Higher the robustness estimates greater is the confidence on the association.

### Causal mediation analysis

We performed causal mediation analysis (VanderWeele, 2016) using the natural effects models for estimation of natural direct (NDE), indirect (NIE) and total causal effects (TCE). The Medflex package (Steen et al., 2017) in R was implemented for the mediation analysis. The causal mediated effects are estimated by fitting the conditional mean models for nested counterfactuals. These models estimate natural direct and indirect effects of an exposure variable on the outcome variable by computing the difference between the potential outcomes with and without mediators. The formulation is given by following equations:

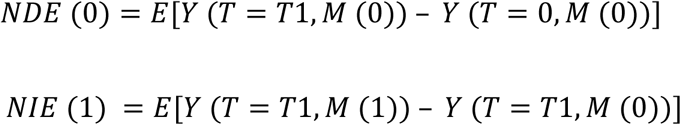

Where, NDE and NIE are the natural direct effects and natural indirect effects (mediated effect), respectively. Y, T and M represents the outcome variable, treatment variable and the mediator variables, respectively. 0 and 1 represent the effects with and without the treatment and mediator variables. The population average natural effects were estimated on the pooled cohort by adjusting for group effects. Inverse probability weighting was used to account for pre-treatment exposure related confounding. The total causal effect is estimated by summing the NDE and NIE.

## Supplementary File

**Figure S1.**
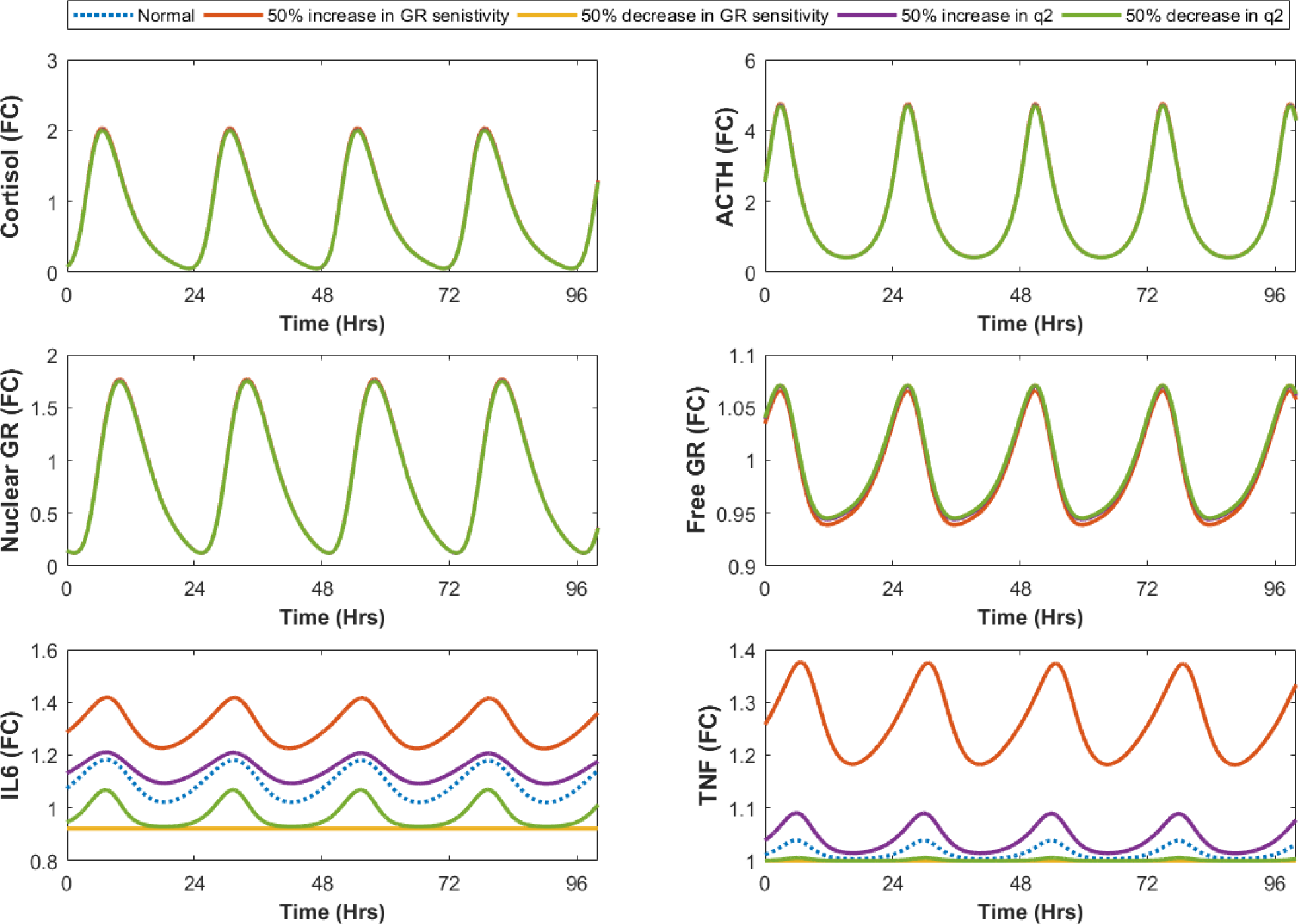
The simulated profiles for varying the GR negative feedback parameters in the immune pathway (GR feedback expressions in Eqn.14). HPA variables were sensitive to increased sensitivity of anti-inflammatory effect of GR’s, which results in elevation of basal pro-inflammatory response and ambient cortisol and ACTH levels.

**Figure S2.**
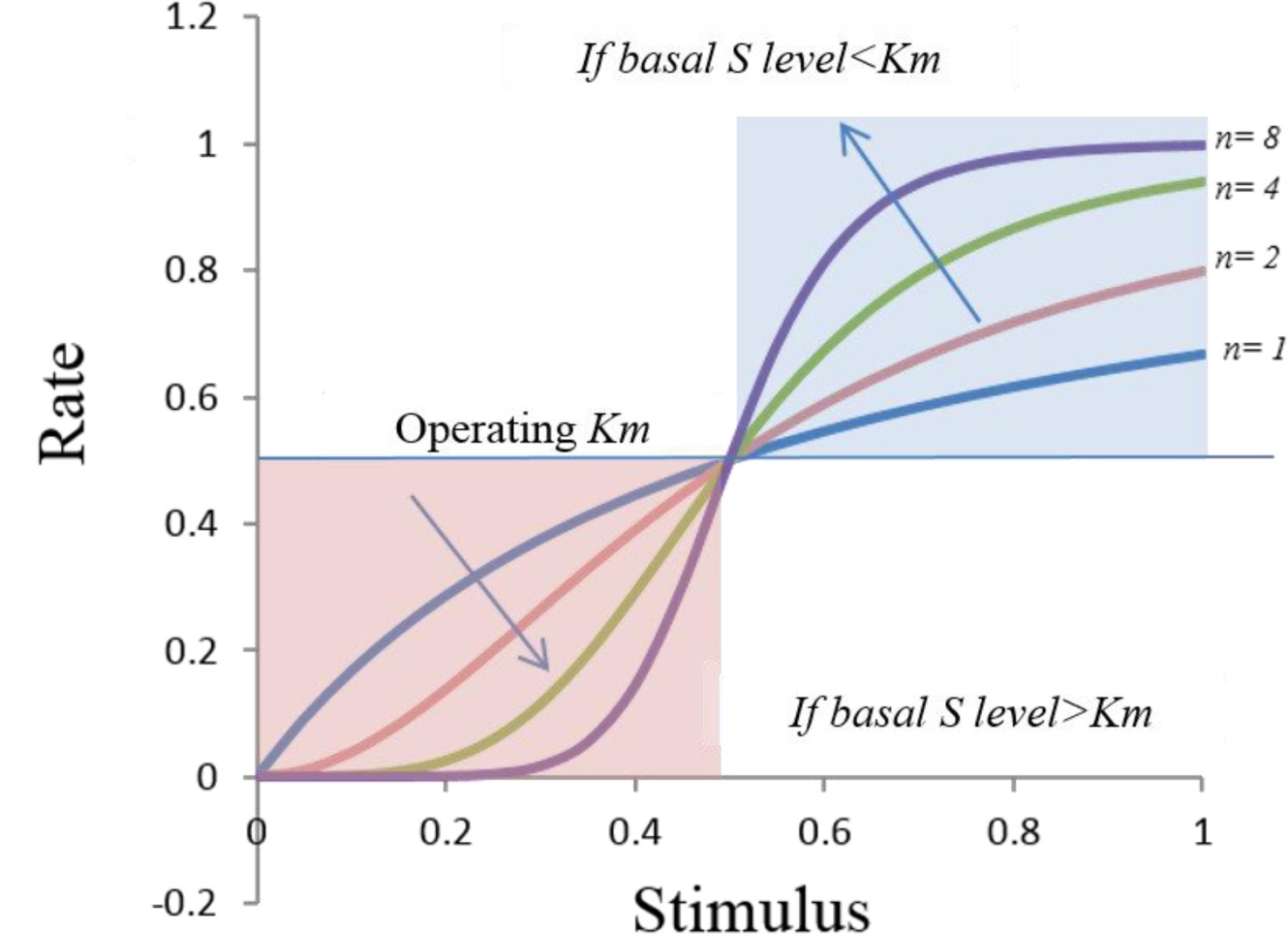
Dose response curves: Simulated representation of change in response rates with respect to changing stimulus (*S*) are shown for different sensitivity (*n*) parameters. The Hill type kinetics as represented as: 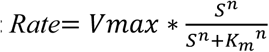 A feedback rate with respect to stimulus (*S)* is determined by three parameters (*Vmax, Km, n*), namely a maximum achievable rate, saturation threshold and sensitivity, respectively (Somvanshi and Venkatesh, 2013). It is observed that the direction of change in response rate is opposite above and below the saturation threshold (*Km*), meaning on increasing the sensitivity the response is lower for the stimulus below the *Km* (red zone) and higher above *Km* (blue zone), whereas, increasing the *Km* reduces the response and vice versa. The inhibitory threshold for anti-inflammatory effect (*q2*) is above the ambient GR levels, therefore increasing GRS decreases the inhibitory effect on increasing GRS (red region) (See Figure 6B(a)). Hence increases pro-inflammatory response due to lower activation of the anti-inflammatory response at relatively lower GR levels as depicted by the sigmoidal dose response curve. The inhibitory threshold for central negative feedback is below the ambient GR levels therefore, increasing GRS increases the inhibitory effect on increasing GRs (blue region) (See Figure 6B(c)) and increases the cortisol suppression effect.

**Table S1.** Demographic and clinical measures of combat veterans with PTSD and controls

| Demographics | PTSD - (N:81) | PTSD + (N:81) | Demographics | PTSD - (N:81) | PTSD + (N:81) |
| --- | --- | --- | --- | --- | --- |
| Sociodemographic |  |  | Medication use (n) |  |  |
| Gender | All males | All males | Sedatives | 4 | 16 |
| Age (Mean±SD) | 32.29 ± 7.60 | 33.20 ± 8.12 | Statins | 1 | 4 |
| Years of Education (Mean±SD) | 14.78 ± 2.32 | 13.81 ± 1.93 | Anti-depressants | 4 | 24 |
| Hispanic/Non-Hispanic (n) | 26/56 | 38/45 | Anticonvulsants | 0 | 9 |
| Smoking (n) | 18 | 33 | Anti-inflammatories | 6 | 9 |
| Alcohol use (Mean±SD) | 1.65 ± 1.08 | 1.38 ± 1.22 | Anti-diabetics | 2 | 2 |
| Biometric measurements (Mean± SD) |  |  | Anti-hypertensives | 5 | 6 |
| BMI | 28.44 ± 4.79 | 30.02 ± 5.047 | Antacids | 3 | 3 |
| Weight | 192.78 ± 35.46 | 205.09 ± 37.77 | Anti-allergics | 4 | 5 |
| Height | 69.05 ± 2.89 | 69.26 ± 2.84 | Pain medicines | 4 | 10 |
| Waist to Hip ratio | 0.88 ± 0.15 | 0.89 ± 0.16 | Comorbid diseases (n) |  |  |
| Pulse | 64.68 ± 11.21 | 72.65 ± 10.33 | Clinical hypertension | 7 | 15 |
| Metabolic measurement (Mean±SD) |  |  | Heart attack | 1 | 1 |
| HbA1c | 5.37 ± 0.44 | 5.39 ± 0.85 | Stable angina | 1 | 3 |
| Cholesterol | 171.35 ± 27.44 | 180.21 ± 35.97 | Diabetics | 2 | 4 |
| LDL | 49.79 ± 13.08 | 47.27 ± 12.05 |  |  |  |
| HDL | 100.56 ± 25.12 | 107.96 ± 32.49 |  |  |  |
| Clinical measures (Mean±SD) |  |  |  |  |  |
| CAPS total current | 9.24 ± 8.39 | 91.55 ± 15.8 |  |  |  |
| CAPS total lifetime | 3.73 ± 5.01 | 69.42 ± 16.91 |  |  |  |
| PCLSCORE | 25.93 ± 8.95 | 61.51 ± 11.84 |  |  |  |
| MCS | 64.34 ± 14.21 | 118.53 ± 19.69 |  |  |  |
| PSQI | 5.19 ± 4.02 | 7.46 ± 5.72 |  |  |  |
| ETISR | 5.98 ± 3.77 | 13.25 ± 3.28 |  |  |  |
| Number of deployment | 1.77 ± 0.89 | 1.82 ± 0.84 |  |  |  |
| MDD diagnosis (n) | 1 | 47 |  |  |  |

**Table S2:** The statistics on the HPA-immune features assessed by Wilcox-ranksum test. The red and yellow color codes for statistical significance of α<=0.5 and 0.1> α >0.05 respectively.

|  | **Controls** |  | **PTSD** |  |  |  |  |  |
| --- | --- | --- | --- | --- | --- | --- | --- | --- |
| **HPA features** | **Mean** | **Std.dev** | **Mean** | **Std. dev** | **p-value** | **q-val** | **Fold change** | **n** |
| Cortisol1 | 13.027 | 5.298 | 14.728 | 6.986 | 0.058 | 0.049 | 1.131 | 162 |
| Cortisol2 | 3.568 | 4.031 | 2.634 | 2.843 | 0.279 | 0.159 | 0.738 | 152 |
| Cortisol difference | 9.392 | 5.048 | 11.524 | 5.406 | 0.008 | 0.013 | 1.227 | 152 |
| Cortisol suppression | 73.856 | 23.004 | 80.613 | 19.967 | 0.037 | 0.037 | 1.091 | 152 |
| ACTH1 | 38.298 | 29.370 | 41.946 | 23.659 | 0.033 | 0.037 | 1.095 | 160 |
| ACTH2 | 15.621 | 12.985 | 14.403 | 11.883 | 0.538 | 0.252 | 0.922 | 152 |
| ACTH difference | 23.208 | 28.861 | 27.164 | 19.460 | 0.005 | 0.013 | 1.170 | 152 |
| ACTH suppression | 55.692 | 26.301 | 64.188 | 28.468 | 0.017 | 0.023 | 1.153 | 152 |
| GR gene methylation | 70.575 | 30.851 | 62.814 | 24.064 | 0.160 | 0.104 | 0.890 | 161 |
| **Immune features** |  |  |  |  |  |  |  |  |
| IC<sub>50</sub>-Dex | 5.304 | 3.356 | 4.361 | 2.556 | 0.108 | 0.075 | 0.822 | 159 |
| IL8 | 10.218 | 6.296 | 11.289 | 12.349 | 0.983 | 0.436 | 1.105 | 162 |
| IL6 | 0.645 | 0.670 | 0.871 | 0.762 | 0.000 | 0.002 | 1.351 | 160 |
| TNFα | 2.751 | 0.789 | 3.966 | 3.870 | 0.007 | 0.013 | 1.442 | 162 |
| IFNγ | 2.076 | 3.355 | 3.260 | 5.131 | 0.102 | 0.075 | 1.570 | 161 |
| IL10 | 1.313 | 1.424 | 1.432 | 1.503 | 0.501 | 0.252 | 1.091 | 162 |
| hs-CRP | 1.638 | 2.390 | 3.362 | 5.205 | 0.007 | 0.013 | 2.052 | 158 |

\*p-values by regression adjusting for non-converted cytosine

**Table S3:**
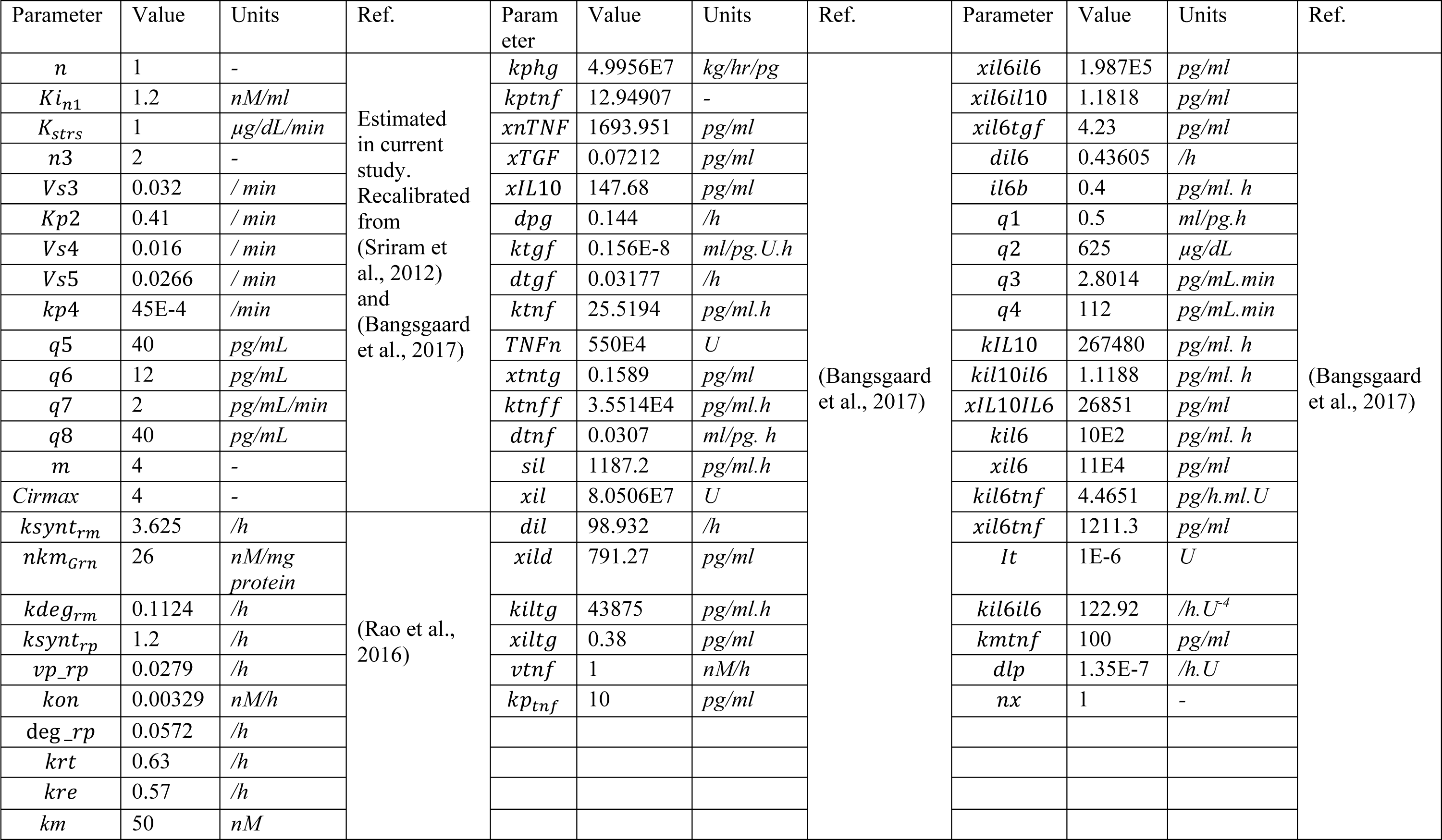
Parameters for the integrated model for the HPA axis, GR signaling and inflammation

### Correlational Analysis

To determine the associations between the HPA axis components and the inflammatory cytokines we performed correlational analysis by estimating Spearman correlational coefficients between HPA axis variables and cytokine in our data. The correlation matrix along with the respective adjusted p-values are reported in Table S1. We observed an expected statistically significant association between cortisol and ACTH related measures (plasma levels and suppression tests). Plasma cortisol levels showed a negative association with GR methylation (ρ=-0.169, p=0.031), IC_50_-DEX (ρ=-0.175, p=0.027), IL6 (ρ=-0.199, p=0.011), IL10 (ρ=-0.222, p=0.0045) and a positive association with IFNγ (ρ=0.175, p=0.026). Cortisol decline showed a negative correlation with IL10 (ρ=-0.172, p=0.034) and positive association with hs-CRP (ρ=-0.15, p=0.068). Cortisol suppression showed strong association with IL6 (ρ=0.189, p=0.019). ACTH levels and cortisol difference also showed a negative trend with IC_50_-DEX. ACTH levels were positively associated with IFNγ(ρ=0.168, p=0.033) and correlated negatively with IL10 (ρ=-0.182, p=0.021). ACTH decline also showed a negative correlation with IL10 (ρ=-0.171, p=0.035) and a positive trend with hs-CRP. ACTH suppression showed a positive association with IL6 (ρ=0.179, p=0.031) and a positive trend with TNFα and hs-CRP. GR methylation was negatively associated with cortisol decline (ρ=-0.161, p=0.049) and positively correlated with IC_50_-DEX (ρ=0.179, p=0.024). IC_50_-DEX showed a strong negative association with IFNγ (ρ=-0.362, p=2.95E-6) and a positive association with IL10 (ρ=0.182, p=0.021).

**Table S4:** The Spearman correlation coefficients and the adjusted p-values for the HPA-immune features.

| HPA -Immune | Cortisol1 |  | Cortisol2 |  | Cortisol difference |  | Cortisol suppression |  | ACTH1 |  | ACTH2 |  | ACTH difference |  | ACTH suppression |  |
| --- | --- | --- | --- | --- | --- | --- | --- | --- | --- | --- | --- | --- | --- | --- | --- | --- |
| features | $\rho$ | p-value | $\rho$ | p-value | $\rho$ | p-value | $\rho$ | p-value | $\rho$ | p-value | $\rho$ | p-value | $\rho$ | p-value | $\rho$ | p-value |
| Cortisol1 | 1 | NA | 0.360 | 5.53E-06 | 0.714 | 0 | 0.017 | 0.83586 | 0.373 | 1.22E-06 | 0.208 | 0.01048 | 0.264 | 0.00112 | 0.090 | 0.2776 |
| Cortisol2 | 0.360 | 5.53E-06 | 1.000 | NA | -0.189 | 0.02074 | -0.896 | 0 | 0.112 | 0.17231 | 0.426 | 6.23E-08 | -0.167 | 0.04117 | -0.385 | 1.58E-06 |
| Cortisol difference | 0.714 | 0 | -0.189 | 0.02074 | 1.000 | NA | 0.571 | 2.38E-14 | 0.225 | 0.0059 | -0.160 | 0.0522 | 0.365 | 4.97E-06 | 0.365 | 6.38E-06 |
| Cortisol suppression | 0.017 | 0.83586 | -0.896 | 0 | 0.571 | 2.38E-14 | 1.000 | NA | 0.005 | 0.94869 | -0.417 | 1.14E-07 | 0.286 | 0.0004 | 0.477 | 1.05E-09 |
| ACTH1 | 0.373 | 1.22E-06 | 0.112 | 0.17231 | 0.225 | 0.0059 | 0.005 | 0.94869 | 1.000 | NA | 0.409 | 2.10E-07 | 0.798 | 0 | 0.249 | 0.00236 |
| ACTH2 | 0.208 | 0.01048 | 0.426 | 6.23E-08 | -0.160 | 0.0522 | -0.417 | 1.14E-07 | 0.409 | 2.10E-07 | 1.000 | NA | -0.140 | 0.08758 | -0.712 | 0 |
| ACTH difference | 0.264 | 0.00112 | -0.167 | 0.04117 | 0.365 | 4.97E-06 | 0.286 | 0.0004 | 0.798 | 0 | -0.140 | 0.08758 | 1.000 | NA | 0.717 | 0 |
| ACTH suppression | 0.090 | 0.2776 | -0.385 | 1.58E-06 | 0.365 | 6.38E-06 | 0.477 | 1.05E-09 | 0.249 | 0.00236 | -0.712 | 0 | 0.717 | 0 | 1.000 | NA |
| GR gene methylation | -0.170 | 0.03139 | -0.126 | 0.12315 | -0.161 | 0.04949 | 0.063 | 0.44217 | -0.104 | 0.19233 | -0.062 | 0.45201 | -0.108 | 0.18806 | -0.082 | 0.32437 |
| IC<sub>50</sub>-DEX | -0.175 | 0.02698 | 0.029 | 0.72387 | -0.140 | 0.09003 | -0.078 | 0.34212 | -0.134 | 0.09535 | 0.039 | 0.6421 | -0.134 | 0.10567 | -0.093 | 0.2672 |
| IL8 | -0.058 | 0.46331 | -0.017 | 0.84007 | -0.116 | 0.1557 | -0.029 | 0.72376 | -0.036 | 0.65574 | 0.070 | 0.39311 | -0.065 | 0.42673 | -0.088 | 0.28789 |
| IL6 | -0.200 | 0.01124 | -0.239 | 0.00327 | -0.001 | 0.98599 | 0.190 | 0.01992 | -0.038 | 0.63638 | -0.181 | 0.02765 | 0.044 | 0.59231 | 0.179 | 0.03092 |
| TNFα | -0.121 | 0.12536 | -0.113 | 0.1654 | -0.089 | 0.27985 | 0.064 | 0.43448 | 0.061 | 0.44461 | -0.118 | 0.15037 | 0.107 | 0.19254 | 0.140 | 0.0897 |
| IFNγ | 0.176 | 0.02595 | 0.079 | 0.33767 | 0.067 | 0.41854 | -0.033 | 0.68919 | 0.168 | 0.03385 | 0.131 | 0.1102 | 0.127 | 0.12272 | -0.005 | 0.95689 |
| IL10 | -0.222 | 0.00454 | -0.097 | 0.23666 | -0.173 | 0.03434 | 0.007 | 0.93564 | -0.182 | 0.02102 | -0.075 | 0.3607 | -0.171 | 0.03587 | -0.044 | 0.59728 |
| hs-CRP | 0.067 | 0.40222 | -0.073 | 0.38039 | 0.151 | 0.06869 | 0.100 | 0.22609 | 0.058 | 0.47194 | -0.096 | 0.25019 | 0.142 | 0.08779 | 0.143 | 0.08757 |
| HPA -Immune | GR gene methylation | | IC<sub>50</sub>-DEX | | IL8 | | IL6 | | TNFα | | IFNγ | | IL10 | | hs-CRP | |
| features | $\rho$ | p-value | $\rho$ | p-value | $\rho$ | p-value | $\rho$ | p-value | $\rho$ | p-value | $\rho$ | p-value | $\rho$ | p-value | $\rho$ | p-value |
| Cortisol1 | -0.170 | 0.03139 | -0.175 | 0.02698 | -0.058 | 0.46331 | -0.200 | 0.01124 | -0.121 | 0.12536 | 0.176 | 0.02595 | -0.222 | 0.00454 | 0.067 | 0.40222 |
| Cortisol2 | -0.126 | 0.12315 | 0.029 | 0.72387 | -0.017 | 0.84007 | -0.239 | 0.00327 | -0.113 | 0.1654 | 0.079 | 0.33767 | -0.097 | 0.23666 | -0.073 | 0.38039 |
| Cortisol difference | -0.161 | 0.04949 | -0.140 | 0.09003 | -0.116 | 0.1557 | -0.001 | 0.98599 | -0.089 | 0.27985 | 0.067 | 0.41854 | -0.173 | 0.03434 | 0.151 | 0.06869 |
| Cortisol suppression | 0.063 | 0.44217 | -0.078 | 0.34212 | -0.029 | 0.72376 | 0.190 | 0.01992 | 0.064 | 0.43448 | -0.033 | 0.68919 | 0.007 | 0.93564 | 0.100 | 0.22609 |
| ACTH1 | -0.104 | 0.19233 | -0.134 | 0.09535 | -0.036 | 0.65574 | -0.038 | 0.63638 | 0.061 | 0.44461 | 0.168 | 0.03385 | -0.182 | 0.02102 | 0.058 | 0.47194 |
| ACTH2 | -0.062 | 0.45201 | 0.039 | 0.6421 | 0.070 | 0.39311 | -0.181 | 0.02765 | -0.118 | 0.15037 | 0.131 | 0.1102 | -0.075 | 0.3607 | -0.096 | 0.25019 |
| ACTH difference | -0.108 | 0.18806 | -0.134 | 0.10567 | -0.065 | 0.42673 | 0.044 | 0.59231 | 0.107 | 0.19254 | 0.127 | 0.12272 | -0.171 | 0.03587 | 0.142 | 0.08779 |
| ACTH suppression | -0.082 | 0.32437 | -0.093 | 0.2672 | -0.088 | 0.28789 | 0.179 | 0.03092 | 0.140 | 0.0897 | -0.005 | 0.95689 | -0.044 | 0.59728 | 0.143 | 0.08757 |
| GR gene methylation | 1.000 | NA | 0.179 | 0.02419 | -0.094 | 0.23525 | -0.019 | 0.81694 | 0.062 | 0.43496 | -0.082 | 0.29996 | 0.037 | 0.64529 | -0.047 | 0.5619 |
| IC<sub>50</sub>-DEX | 0.179 | 0.02419 | 1.000 | NA | 0.012 | 0.88296 | 0.122 | 0.12727 | 0.027 | 0.73594 | -0.362 | 2.95E-06 | 0.182 | 0.02134 | -0.059 | 0.46957 |
| IL8 | -0.094 | 0.23525 | 0.012 | 0.88296 | 1.000 | NA | 0.105 | 0.18532 | 0.315 | 4.38E-05 | 0.026 | 0.74667 | 0.209 | 0.00751 | -0.099 | 0.21564 |
| IL6 | -0.019 | 0.81694 | 0.122 | 0.12727 | 0.105 | 0.18532 | 1.000 | NA | 0.471 | 3.27E-10 | -0.206 | 0.00921 | 0.508 | 7.04E-12 | 0.455 | 2.52E-09 |
| TNFα | 0.062 | 0.43496 | 0.027 | 0.73594 | 0.315 | 4.38E-05 | 0.471 | 3.27E-10 | 1.000 | NA | -0.033 | 0.67673 | 0.534 | 2.65E-13 | 0.139 | 0.08209 |
| IFNγ | -0.082 | 0.29996 | -0.362 | 2.95E-06 | 0.026 | 0.74667 | -0.206 | 0.00921 | -0.033 | 0.67673 | 1.000 | NA | -0.521 | 1.39E-12 | 0.071 | 0.37469 |
| IL10 | 0.037 | 0.64529 | 0.182 | 0.02134 | 0.209 | 0.00751 | 0.508 | 7.04E-12 | 0.534 | 2.65E-13 | -0.521 | 1.39E-12 | 1.000 | NA | 0.067 | 0.39946 |
| hs-CRP | -0.047 | 0.5619 | -0.059 | 0.46957 | -0.099 | 0.21564 | 0.455 | 2.52E-09 | 0.139 | 0.08209 | 0.071 | 0.37469 | 0.067 | 0.39946 | 1.000 | NA |

**Table S5.**
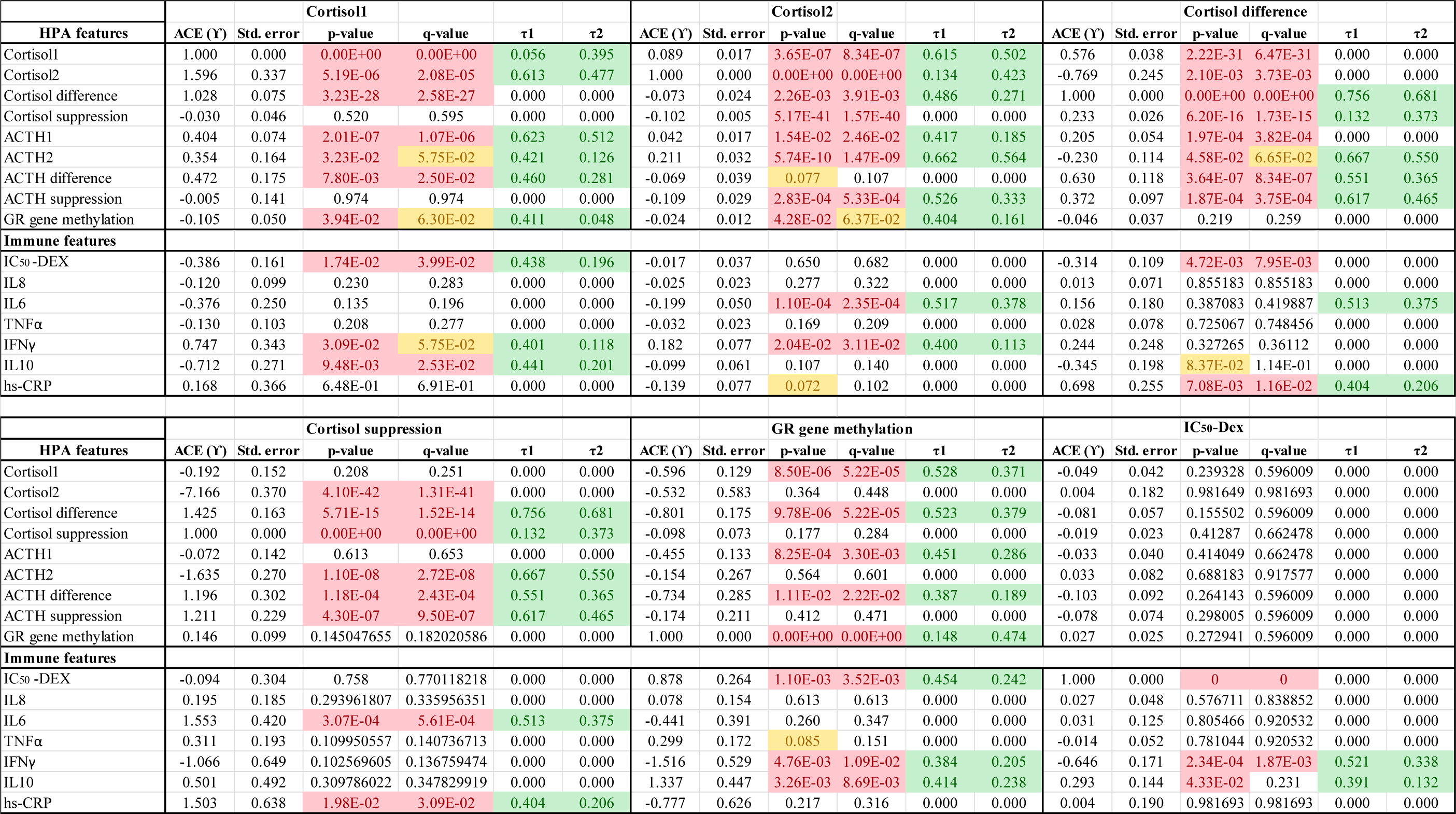
Population level average causal effects. Robustness estimate τ1 representing the coefficient of unobserved confounder at which ACE=0, robustness estimate τ2 representing the coefficient of unobserved confounder at which ACE becomes statistically insignificant (p>0.05). Red, yellow and green shade highlights the p and q values <0.05, between 0.05 and 0.1 and τ_2_ >0.2, respectively.

**Table S6.** Estimates and statistics for natural direct, natural indirect and total causal effect of glucocorticoid receptor methylation mediated through measures of glucocorticoid receptor sensitivity (Post dexamethasone cortisol levels and IC_50_-dex lysozyme suppression) on cytokines. Red and yellow shade highlights the p and q values <0.05 and between 0.05 and 0.1, respectively.

| Cytokines | Natural Direct Effect |  |  |  |  |  | Natural Indirect Effect |  |  |  |  |  | Total Causal Effect |  |  |  |  |  |
| --- | --- | --- | --- | --- | --- | --- | --- | --- | --- | --- | --- | --- | --- | --- | --- | --- | --- | --- |
| Post-Dex cortisol related mediated effects |  |  |  |  |  |  |  |  |  |  |  |  |  |  |  |  |  |  |
| | NDE<br>($\psi_d$) | Std.<br>error | CI-lower | CI-upper | p-value | q-value | NIE<br>($\psi_i$) | Std.<br>error | CI-lower | CI-upper | p-value | q-value | TCE<br>($\psi_t$) | Std.<br>error | CI-lower | CI-upper | p-value | q-value |
| IL8 | 0.052 | 0.149 | -0.240 | 0.344 | 0.728 | 0.728 | 0.040 | 0.035 | -0.028 | 0.109 | 0.251 | 0.251 | 0.092 | 0.145 | -0.191 | 0.375 | 0.524 | 0.629 |
| IL6 | -0.129 | 0.352 | -0.819 | 0.561 | 0.714 | 0.728 | 0.229 | 0.100 | 0.033 | 0.424 | 0.022 | 0.130 | 0.100 | 0.349 | -0.584 | 0.783 | 0.775 | 0.775 |
| TNFα | 0.152 | 0.190 | -0.220 | 0.524 | 0.422 | 0.633 | 0.058 | 0.034 | -0.009 | 0.125 | 0.092 | 0.193 | 0.210 | 0.188 | -0.159 | 0.579 | 0.265 | 0.398 |
| IFNγ | -1.196 | 0.521 | -2.217 | -0.175 | 0.022 | 0.088 | -0.147 | 0.097 | -0.336 | 0.043 | 0.129 | 0.193 | -1.343 | 0.529 | -2.379 | -0.307 | 0.011 | 0.047 |
| IL10 | 0.933 | 0.428 | 0.094 | 1.771 | 0.029 | 0.088 | 0.096 | 0.076 | -0.054 | 0.246 | 0.208 | 0.250 | 1.029 | 0.426 | 0.194 | 1.864 | 0.016 | 0.047 |
| hs-CRP | -1.130 | 0.618 | -2.341 | 0.081 | 0.067 | 0.135 | 0.163 | 0.098 | -0.030 | 0.355 | 0.097 | 0.193 | -0.967 | 0.607 | -2.157 | 0.222 | 0.111 | 0.222 |
| IC<sub>50</sub>-Dex lysozyme suppression related mediated effects |  |  |  |  |  |  |  |  |  |  |  |  |  |  |  |  |  |  |
| | NDE<br>($\psi_d$) | Std.<br>error | CI-lower | CI-upper | p-value | q-value | NIE<br>($\psi_i$) | Std.<br>error | CI-lower | CI-upper | p-value | q-value | TCE<br>($\psi_t$) | Std.<br>error | CI-lower | CI-upper | p-value | q-value |
| IL8 | 0.080 | 0.147 | -0.207 | 0.368 | 0.584 | 0.701 | 0.015 | 0.028 | -0.040 | 0.069 | 0.598 | 0.718 | 0.095 | 0.144 | -0.188 | 0.378 | 0.511 | 0.613 |
| IL6 | 0.063 | 0.355 | -0.634 | 0.759 | 0.859 | 0.859 | 0.059 | 0.066 | -0.069 | 0.188 | 0.365 | 0.718 | 0.122 | 0.354 | -0.572 | 0.816 | 0.730 | 0.730 |
| TNFα | 0.220 | 0.192 | -0.157 | 0.597 | 0.253 | 0.379 | 0.000 | 0.028 | -0.055 | 0.055 | 0.999 | 0.999 | 0.220 | 0.189 | -0.150 | 0.590 | 0.244 | 0.366 |
| IFNγ | -0.946 | 0.536 | -1.997 | 0.105 | 0.078 | 0.176 | -0.327 | 0.123 | -0.568 | -0.086 | 0.008 | 0.046 | -1.273 | 0.525 | -2.302 | -0.244 | 0.015 | 0.056 |
| IL10 | 0.856 | 0.423 | 0.027 | 1.684 | 0.043 | 0.176 | 0.147 | 0.086 | -0.021 | 0.315 | 0.086 | 0.258 | 1.003 | 0.426 | 0.168 | 1.837 | 0.019 | 0.056 |
| hs-CRP | -1.006 | 0.589 | -2.161 | 0.149 | 0.088 | 0.176 | 0.052 | 0.098 | -0.140 | 0.244 | 0.596 | 0.718 | -0.954 | 0.602 | -2.134 | 0.226 | 0.113 | 0.226 |

